# A cryptic pocket allosterically modulates oligosaccharide binding to DC-SIGN

**DOI:** 10.1101/2025.08.19.671048

**Authors:** Jonathan Lefèbre, Maurice Besch, Marcelo Daniel Gamarra, Jan-Oliver Kapp-Joswig, Annika Balke, Henry Flatau, Gregor Suchy, Elena Georgieva, Patrick Scheerer, Bettina G. Keller, Carlos Pablo Modenutti, Christoph Rademacher

## Abstract

DC-SIGN is a C-type lectin receptor expressed on antigen-presenting cells, crucial for pathogen recognition and immune modulation. Here, we identify and characterize a previously unrecognized cryptic allosteric pocket in DC-SIGN using molecular dynamics simulations, NMR spectroscopy, cryogenic electron microscopy and biochemical assays. Rotation of the gatekeeper residue M270 exposes the pocket whose occupancy modulates glycan binding. Mutations M270F and T314A mimic the occupied and unoccupied states of this pocket, respectively, shifting the conformational equilibrium of α-helix 2 and altering oligosaccharide affinity via the extended carbohydrate binding site. While Ca²⁺ coordination at the canonical binding site remains unaffected, our data reveal a complex interplay between the Ca²⁺ binding sites and the canonical and extended glycan binding surfaces. These findings uncover a hierarchical allosteric mechanism that enables selective tuning of glycan affinity and suggest the cryptic pocket as a novel target for drug discovery in C-type lectins.

## Introduction

Dendritic Cell-Specific Intercellular adhesion molecule-3-Grabbing Non-integrin (DC-SIGN) is a Ca²⁺-dependent glycan-binding C-type lectin receptor expressed on antigen-presenting cells, including dendritic cells and macrophages (*1, 2*). It plays a central role in bridging innate and adaptive immunity by discriminating self from non-self, facilitating pathogen recognition, cellular adhesion and immunological synapse formation (*3–9*). The receptor recognizes a broad range of carbohydrate structures, most prominently high-mannose and fucose-containing glycans, such as Lewis X (*10–12*). These include glycans from bacterial and fungal pathogens, as well as viral glycoproteins exploited, for example, by HIV-1 and SARS-CoV-2 for cellular entry and dissemination (*13–19*). DC-SIGN also recognizes self-antigens on endogenous glycoproteins, underscoring the remarkable diversity of ligands and distinct biological outcomes, ranging from pathogen uptake and immune activation to tolerance (*1, 20, 21*). Consequently, DC-SIGN not only orchestrates immune surveillance but is also hijacked by pathogens to promote infection, making it an attractive target for antiviral therapeutics, immunomodulatory interventions, and targeted delivery to immune cells (*9, 22–24*).

Critical for the function of DC-SIGN is its ability to recognize a diverse repertoire of glycan structures through its carbohydrate recognition domain (CRD). The DC-SIGN CRD adopts a typical C-type lectin domain fold and binds three Ca²⁺ ions at its long loop region, one of which is coordinated by the conserved Glu-Pro-Asn (EPN) motif, essential for recognizing mannose- and fucose-containing glycans at the canonical carbohydrate binding site (*25, 26*). To achieve functional binding, DC-SIGN engages not only this canonical binding site but also an extended carbohydrate binding site formed by β-strands 2-4, ⍺-helix 2 and residues of the ⍺2-β2 connecting loop (Fig. 1A) (*25*). This extended site supports recognition of oligosaccharides in multiple binding modes and contributes secondary contacts that enhance binding affinity and specificity (*27*).

**Figure 1.**
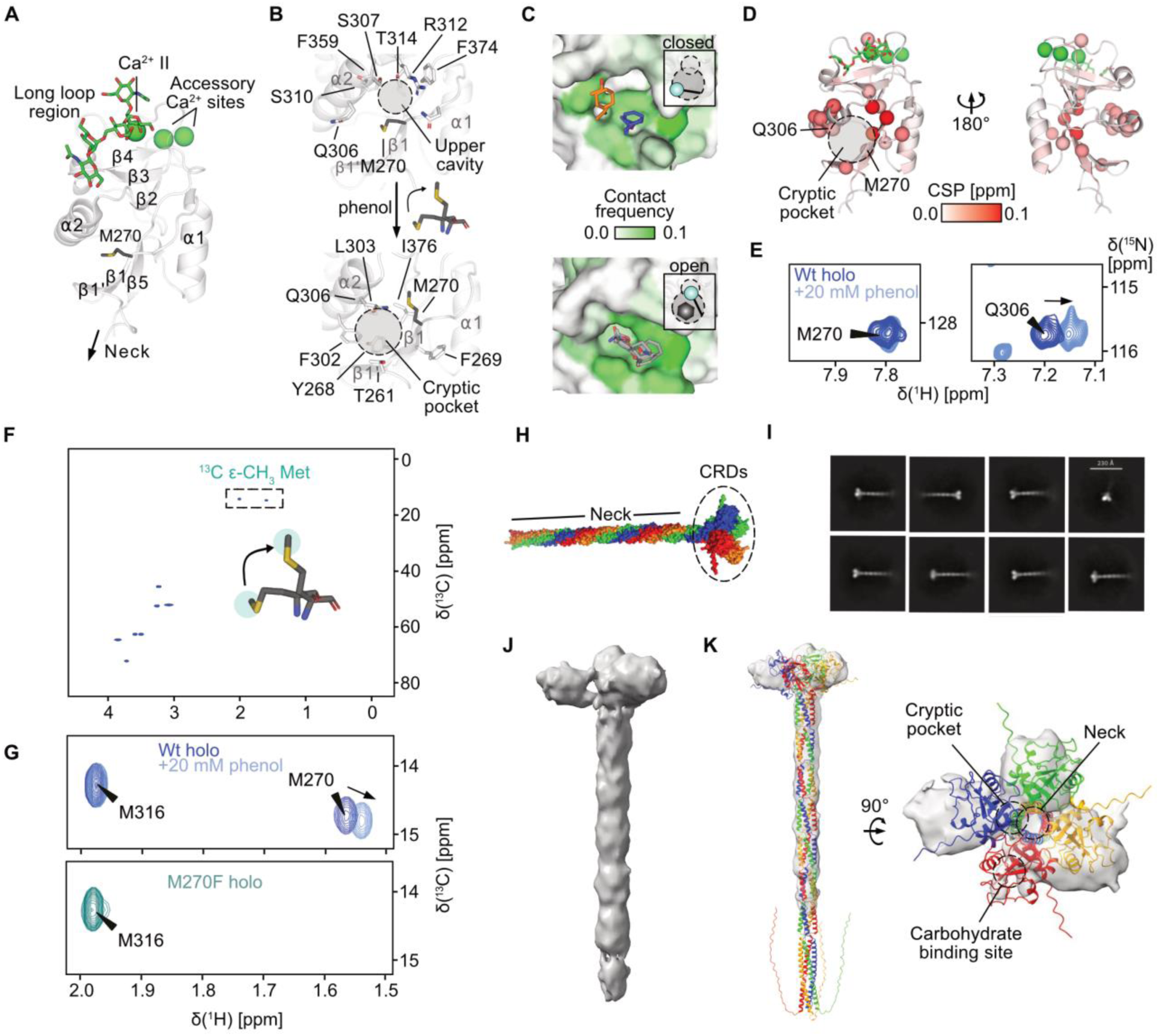
DC-SIGN harbors a cryptic secondary site. (**A**) X-ray crystallographic structure of DC-SIGN CRD in complex with GlcNAc_2_Man_3_ (PDB ID: 1K9I) with secondary structure numbering (*26*). The glycan interacts with the canonical carbohydrate binding site via Ca²⁺ II complexed in the long loop region and an extended carbohydrate binding site formed by β-strands 2-4, ⍺-helix 2 and a short loop connecting ⍺2 and β2. A previously identified secondary site between ⍺-helix 2 and β-strands 1’, 1 and 5 is centered by residue M270. (**B**) Mixed-solvent MD simulations with phenol reveal that upon binding, M270 rotates into an upper cavity lined by residues Q306, S307, S310, R312, T314, F374, F359 and N272. This opens a hydrophobic cryptic pocket formed by residues T261, Y268, F269, M270, F302, L303 and I376. (**C**) FTMap analysis of the closed pocket (top) indicated only two smaller solvent probe clusters (orange and blue sticks) and high contact frequencies adjacent to the cryptic site. The open cryptic pocket (bottom) revealed one larger cluster (grey sticks) that extends into the pocket suggesting higher druggability than the closed pocket (*51*). (**D**) CSPs observed in ¹H-¹⁵N HSQC NMR experiments upon addition of 20 mM phenol to holo DC-SIGN CRD validate the binding site of phenol at the cryptic pocket. C⍺ atoms of residues experiencing a CSP > 0.025 ppm are shown as spheres. (**E**) Resonances of residues M270 and Q306 are shown. While Q306 at the cryptic site shows a large CSP upon phenol binding, a small perturbation for M270 suggests limited changes in the amide backbone of the residue. (**F**) Labeling of DC-SIGN CRD wildtype with ^13^C Met provides ¹H-^13^C HSQC NMR spectra of the two methionine sidechains in residues M316 and M270. (**G**) The setup enables direct observation of phenol opening the cryptic pocket, as indicated by a CSP induced in the M270 resonance (top). Assignment of the methionine resonances was done using a M270F mutant (bottom). (**H**) Alphafold3 model of the DC-SIGN ECD forming tetramers under physiological conditions via the neck domain (Fig. S5). (**I**) Representative 2D class averages of DC-SIGN ECD particles, showing predominantly side and intermediate views, with top views being underrepresented. Scale bar: 230 Å. (**J**) Cryo-EM map (colored in grey) of DC-SIGN ECD showing the expected tetrameric topology in absence of a glycan ligand, with three of the four CRDs visibly resolved. (**K**) Rigid body docking of the cryo-EM map to the Alphafold3 model reveals a different arrangement of the CRDs, likely resulting from high flexibility of the CRD-neck junction. The relative position of the cryptic pocket, the carbohydrate binding site and the CRD-neck junction are highlighted with dotted circles.

Under physiological conditions the DC-SIGN extracellular domain (ECD) forms tetramers via its neck region allowing for multivalent recognition of glycans, further amplifying ligand binding by avidity effects, eventually leading to endocytosis and immune signaling (*28–30*). The cellular response is highly context-dependent and can vary significantly with antigen structure and the glycan involved (*31–34*). Accordingly, while this multilevel architecture enables functional plasticity, it also raises questions about how glycan binding at the CRD translates into cellular responses, particularly in light of the structure–function relationship that governs signaling and endocytic activity of other C-type lectin receptors, such as langerin, MGL and CLEC5A (*35–38*). Importantly, understanding conformational dynamics and regulatory mechanisms may enable the rational design of compounds that inhibit, activate or fine-tune the activity of DC-SIGN in the context of immunity and infection.

Despite its therapeutic potential, designing small molecules that directly target the carbohydrate binding site of DC-SIGN with high selectivity and affinity remains challenging. The site is shallow, polar, and binding usually involves a large entropic penalty due to loop flexibility and solvent exposure resulting in low-affinity and high promiscuity of DC-SIGN-carbohydrate interactions (*39, 40*). An emerging strategy to circumvent these limitations involves targeting secondary sites that are distinct from the canonical carbohydrate binding site (*41*). In many C-type lectins, including DC-SIGN, structural and computational studies have revealed the presence of such sites, some of which are proposed to modulate glycan recognition allosterically (*42–48*). Compared to the canonical carbohydrate binding site, these secondary sites can provide increased selectivity, functional specificity, and new opportunities for therapeutic intervention. Fragment-based drug discovery has been instrumental in identifying such secondary sites in C-type lectins and in DC-SIGN. Several fragment hits have been reported to bind to previously unrecognized pockets, including one distal site between ⍺-helix 2 and β-strands 1’, 1 and 5 that is centered by residue M270 (Fig. 1A). The same site was later used to target DC-SIGN-expressing cells using heteromultivalent liposomes and was suggested to allosterically modulate Ca²⁺-dependent carbohydrate binding by an unknown mechanism (*49*).

In this study, we combined molecular dynamics (MD) simulations, nuclear magnetic resonance (NMR) spectroscopy and biochemical assays to identify and characterize a previously unrecognized flexible druggable pocket in DC-SIGN, accessible only upon conformational rearrangement. We show that the rotational shift of the “gatekeeper” residue methionine M270 exposes this cryptic pocket whose occupancy modulates glycan binding allosterically at the carbohydrate binding site. Introducing M270F and T314A mutations mimics the occupied and the unoccupied states of the pocket, respectively, and shifts the conformational equilibrium of ⍺-helix 2, thereby tuning oligosaccharide affinity via the extended carbohydrate binding site. We excluded a direct effect on the Ca²⁺ cage but find evidence for a more complex interplay of the Ca²⁺ sites, the canonical and the extended carbohydrate binding site. Together, these results reveal a hierarchical allosteric mechanism that primes DC-SIGN for selective glycan engagement and opens new avenues for drug discovery targeted at C-type lectin receptors.

## Results

### Gatekeeper residue M270 controls cryptic pocket opening

We have previously used *in silico* methods and fragment screening to explore druggability and allosteric potential of secondary sites in DC-SIGN (*44, 45, 49*). While several fragment hits could be identified, *in silico* druggability as evaluated from X-ray crystallographic structures was low. To expand our knowledge on the local dynamics and druggability of the fragment sites we conducted mixed solvent MD simulations with the DC-SIGN CRD in the presence or absence of phenol (*50*). Simulations in 5% v/v water/phenol mixtures identified several hotspots corresponding to previously identified secondary fragment binding sites remote of the carbohydrate binding site of DC-SIGN (Fig. S1A). While backbone conformational changes where limited throughout the simulations, one of the hotspots located between ⍺-helix 2 and β-strand 5 was only revealed upon rotation of the thioether sidechain of residue M270 into a small upper cavity lined by residues of the C-terminal end of ⍺-helix 2, β-strand 2 and the short ⍺2-β2 loop connecting them (Fig. 1B). Upon movement of M270, water is released from the upper cavity, and a hydrophobic cryptic pocket is exposed (Fig. S1B). Analysis of this open state by FTMap suggested one larger consensus site, extending into the opened hydrophobic pocket (Fig. 1C) (*51*). Although the open state was significantly less populated in the absence of phenol, a population of 34 % indicated an equilibrium between the open and closed state in the absence of phenol and conformational selection of the open state by small molecules (Fig. S1C, D).

To experimentally verify the interaction of small organic molecules with the cryptic pocket we titrated phenol at high excess to uniformly ¹⁵N-labeled DC-SIGN CRD in ¹H-¹⁵N heteronuclear single quantum coherence (HSQC) NMR experiments. Compared to the sample without phenol, we observed strong chemical shift perturbations (CSPs) in residues of the predicted cryptic pocket as well as the upper cavity, to which the M270 side chain rotates to. Smaller CSPs were observed for more distant sites including residues in the adjacent ⍺-helix 2 and the long loop region (Fig. 1D). To further test direct interaction with the pocket, we repeated the titration with a previously established DC-SIGN CRD M270F mutant that blocked secondary site binding (*49*). We observed significantly less CSPs around that site, validating this approach (Fig. S2).

Although ¹H-¹⁵N HSQC NMR allowed us to infer direct binding to the site around M270, CSPs in the wildtype M270 resonance were small, indicating the backbone ¹⁵N amide to experience little change in its chemical environment (Fig. 1E). Since the mechanism of opening of the pocket largely depends on rotation of the methionine sidechain instead of backbone movements, we reasoned that isotopic ¹³C labeling of the terminal CH_3_ group of the thioether side chain of M270 with ^13^C ε-methyl methionine (^13^C Met) could serve as a more sensitive indicator for cryptic pocket opening. We expressed ^13^C Met-labeled DC-SIGN CRD wildtype and assigned the corresponding ¹H-¹³C resonance of M270 by comparison with the ¹H-¹³C HSQC NMR spectra of ¹³C-Met-labeled DC-SIGN CRD M270F mutant (Fig. 1F, G and Fig. S3). In line with our observations from mixed solvent MD simulations, addition of phenol induced a CSP in the M270 resonance, suggesting the ^13^C Met sidechain to experience a change in its chemical environment. In contrast, the other methionine resonance corresponding to M316, remained unchanged (Fig. 1G).

While a co-crystal structure harboring a small molecule in the cryptic pocket is not available, a previous X-ray crystallographic structure of DC-SIGN harbors a tryptophan side chain from a symmetry mate protein of the unit cell in the cryptic pocket (PDB ID: 1SL5) (*52*). Here, the open state is additionally stabilized by a predicted hydrogen bond of the M270 thioether sidechain to adjacent residue Q306 (Fig. S4). Yet, considering the DC-SIGN ECD to form homotetramers under physiological conditions, the spatial arrangement of the CRDs might affect accessibility of the secondary site in the oligomer (Fig. 1H) (*30*).

To approximate the relative orientation of the CRDs in the ECD tetramer, we conducted cryogenic electron microscopy (cryo-EM) experiments with DC-SIGN ECD (Fig. S6). Although the resulting cryo-EM maps had low global resolution we clearly observed the expected topology of the tetramer (Figs. 1I, J and S7). While the first repeats of the coiled-coiled neck domain were rigid, the relative CRD orientation appeared to be highly variable, where only three of four CRDs were visibly resolved. This is consistent with previous reports on the flexibility of the CRD-neck linkage and suggested exposure of the secondary site in the ECD tetramer (Fig. 1K) (*53*). To further prove the accessibility of the secondary site in the ECD in solution, we expressed ^13^C Met-labeled DC-SIGN ECD wildtype and recorded ¹H-^13^C transverse relaxation-optimized spectroscopy (TROSY) NMR spectra in the presence and absence of 20 mM phenol. In line with our previous observation with the CRD, the M270 resonance experienced a CSP in the presence of phenol (Fig. S8). As the CSP trajectory resembled that of the CRD spectra, we concluded that the cryptic site is accessible in the ECD tetramer.

Taken together, these results demonstrate that binding of phenol to the previously described secondary site underlies opening of a cryptic pocket due to rotational shift of the thioether side chain of the gatekeeper residue M270. This mechanism is preserved in the tetrameric ECD suggesting relevance of the site under physiological conditions.

### Residues of the secondary site are hubs in a Ca²⁺-responsive allosteric network

We have previously hypothesized that binding to the secondary site centered by M270 could allosterically modulate Ca²⁺-dependent carbohydrate binding of DC-SIGN (*49*). Our mixed solvent MD simulations and HSQC NMR experiments with phenol suggested a change in local dynamics upon binding of small molecules to the secondary site but showed only minor long-range effects on the primary Ca²⁺-dependent carbohydrate binding site, likely due to low affinity of phenol. As a major hallmark of allosteric modulation is reciprocity, changes or perturbations at the Ca²⁺-dependent carbohydrate binding site should propagate to the secondary site leading to significant changes in HSQC NMR experiments (*54*).

To address the coupling between the secondary site and the carbohydrate and Ca²⁺ binding sites, we recorded ¹H-¹³C HSQC NMR spectra of ^13^C Met-labeled DC-SIGN CRD in the absence (apo) and presence (holo) of Ca^2+^, as well as under mannose-bound conditions. While mannose had no detectable effect, removal of Ca²⁺ induced CSPs and line broadening specifically in the M270 sidechain resonance but not in M316, indicating conformational or dynamic changes that originate at the Ca²⁺ site and extend to the secondary site (Fig. S9A, B). Notably, the CSP direction upon Ca²⁺ removal was opposite to that induced by phenol, and phenol addition to the apo protein shifted the signal back towards the holo state, resulting in a linear correlation of chemical shifts and supporting a model of conformational coupling between the Ca²⁺ sites and the cryptic pocket (Fig. S9C) (*55*). Complementing these observations, ¹H-¹⁵N HSQC NMR titrations with CaCl₂ revealed widespread CSPs and altered exchange regimes across the CRD, particularly in α-helix 2 and β-strands 2–4, underscoring global structural rearrangements that link the cryptic pocket and extended glycan binding site through a Ca²⁺-responsive allosteric network (Fig. S9D-F).

To gain deeper insight into how an allosteric signal could be communicated between the Ca²⁺ and the secondary site, we performed microsecond-scale all-atom MD simulations of both the apo and holo states of the DC-SIGN CRD in explicit water. We analyzed the normalized mutual information (NMI) between dihedral angles for all pairs of residues, which measures the degree of conformational coupling, therefore revealing networks of communicating residues (*38, 56*). The NMI graphs for the apo and holo states of DC-SIGN revealed complex, layered networks of both backbone dihedral angles (φ-ψ) and side-chain dihedral angles (χ) correlations across the whole protein structure. While the apo state exhibited a more fragmented network, the holo state showed increased connectivity, especially within regions containing β-strands 2, 3, and 4 and α-helix 2 (Figs. 2A and S10). This was further confirmed by comparing the node degrees as an indicator of connectivity or importance each residue in the NMI networks. High node degree emerged in a broader set of residues under holo conditions, with the exception of a few long-loop residues that showed stronger connectivity in the apo state, similar to observations in the related CTL langerin (Fig. 2B, E) (*38*). In particular, residues in β-strands 2, 3 and 4, ⍺-helix 2 and the secondary site exhibited high node degree and acted as network hubs in the holo state but drastically lost importance in the apo state (Fig. 2C, D). Furthermore, analyzing hydrogen bond populations along the MD trajectories revealed significant changes in the same regions of the CRD, indicating that correlated conformations in the NMI network are underlying physical interactions (Table S1). Consistent with our observations from experimental CaCl_2_ titrations, these differences suggested Ca²⁺ binding to enhance both local and distant residue interactions. In particular for residues lining the extended carbohydrate binding site and the upper cavity of the secondary site in β-strands 2, 3, 4 and ⍺-helix 2, a high node degree coincided with conformational changes observed in our ¹H-¹⁵N HSQC NMR CaCl_2_ titration experiments.

**Figure 2.**
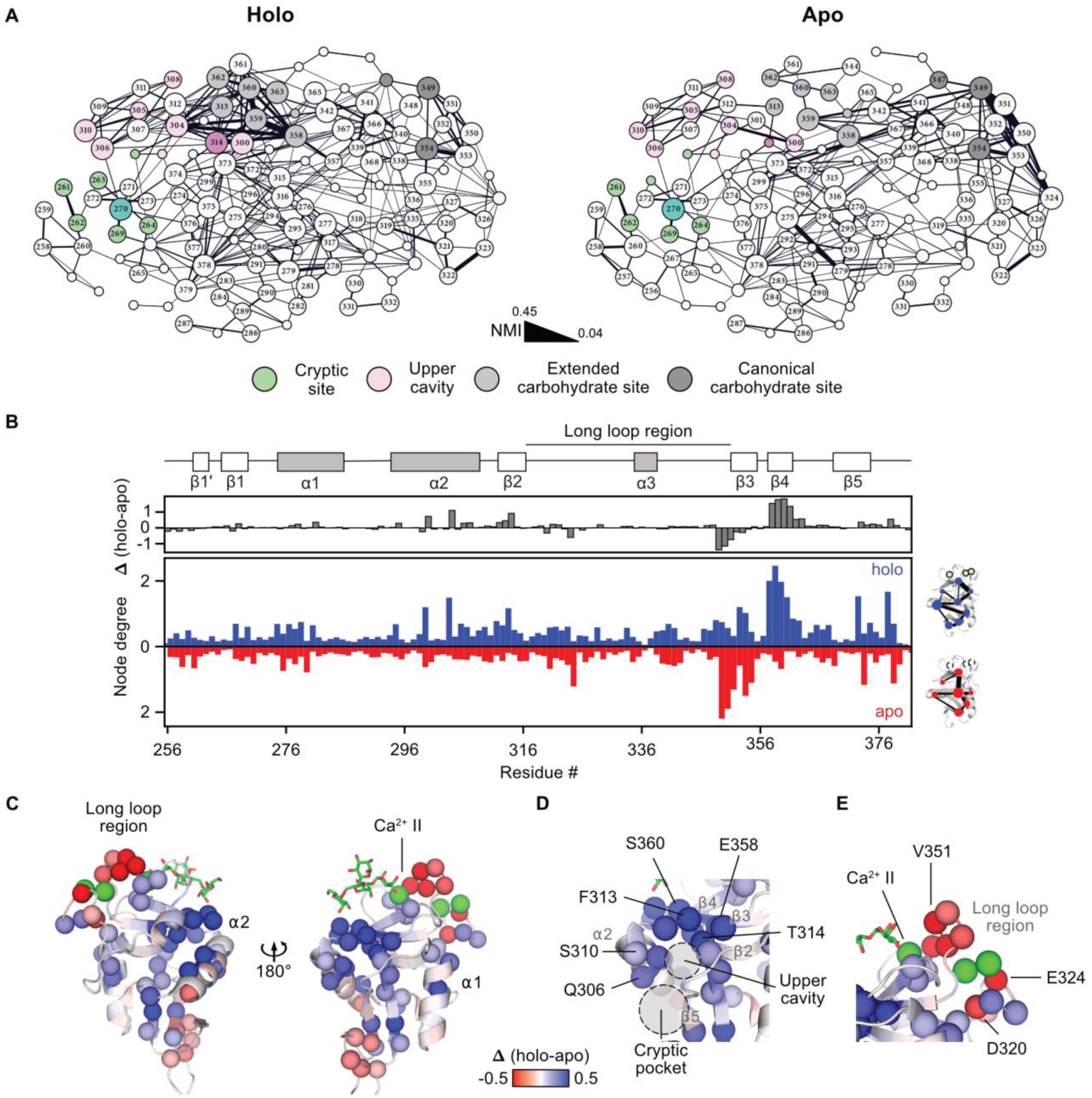
MD simulations and NMI analysis of apo and holo DC-SIGN reveal network hubs. (**A**) Network representation of the NMI graphs computed for the holo (left) and apo (right) state of DC-SIGN CRD. Edge thickness corresponds to computed NMI values (NMI threshold 0.04). Nodes exceeding a degree of 5 are defined as hubs and additionally as highlights if at least one edge exceeds an NMI value of 0.1. Nodes are scaled based on their importance in the network: Highlight and hub > highlight > hub > none. Nodes are colored according to their functional role in the CRD structure. Residues of the cryptic site and the upper cavity are shown in green and pink, respectively. Residues of the canonical and the extended carbohydrate binding site are shown in gray and light gray, respectively. A clear shift of connectivity towards the extended carbohydrate binding site is observed, when comparing the holo to the apo network (Fig. S10). (**B**) Calculation of edge weighted node degrees from NMI analysis reveals residues with high importance in the holo (blue) and the apo (red) network of the CRD. The Δ node degree (holo – apo node degree) illustrates the importance of each residue in the holo network. (**C**) and (**D**) Mapping of Δ node degree on the CRD structure reveals clustering of holo network hubs at the extended carbohydrate site in β2-4 (E358, F359, S360, G361, N362, G363, W364) and the upper cavity (T314, Q300, Q304, Q306, S307, S310). (**E**) In contrast, negative Δ node degrees in residues of the long loop region, indicate higher connectivity in the apo state.

### Mutation of hub residues alter glycan binding properties of DC-SIGN

Overall, our data suggested a Ca²⁺-responsive network in DC-SIGN CRD potentially responsible for allosteric signal transmission between the Ca²⁺ sites and the cryptic pocket. In this network, residues in ⍺-helix 2, the conserved hydrophobic core and β-strands 3 and 4 involved in the formation of the extended carbohydrate binding site appeared to take a central role in the network. We reasoned that mutation of NMI hubs and the cryptic pocket should lead to perturbations of the network, eventually translating into chemical shift changes in ¹H-¹⁵N HSQC NMR spectra (*54*).

To test our hypothesis, we initially selected eight point mutations based on insights from our MD simulations and NMR experiments and expressed their ¹⁵N-labeled CRDs. All mutant holo CRDs produced well-dispersed spectra indicating correct folding of the proteins (Fig. S12). Superposition of the mutant to the wildtype spectra revealed large CSPs in residues in close proximity to and over 10 Å away from the mutation sites. These remote, non-nearest neighbor effects were most pronounced for the T314A, E358A, F359L and W364F mutants, suggesting mutation of hub residues to globally affect the CRD fold (Fig. S13). In mannose titrations, all mutant CRDs showed characteristic CSPs upon addition of mannose, further confirming correct folding and activity of the proteins (Fig. S14). Nevertheless, fitting of dissociation constants (K_D_s) revealed significant reduction in affinity for mutations E358A and F359L and to a lesser extent for mutants F313A and W364F at the extended carbohydrate binding site (Figs. S15 and S16).

To avoid direct interference with the canonical carbohydrate binding site and Ca²⁺ cage in subsequent assays, we selected T314A, a key hub residue identified in MD simulations that induced long-range CSPs while only marginally affecting mannose affinity. As T314 is located in the upper cavity where the M270 side chain rotates to upon opening of the cryptic pocket, it lays central between other hubs of the NMI network located in ⍺-helix 2, the conserved hydrophobic core and β-strands 3 and 4 involved in the formation of the extended carbohydrate binding site, suggesting this residue to also act as a major hub in allosteric signal transmission from the secondary site. Although less prominent in the NMI analysis, we also selected the mutation M270F of the cryptic pocket gatekeeper residue M270 for experimental validation. As it mimics the cryptic pocket to be occupied by a small-molecule ligand similar to phenol, we reasoned that this could allow us to evaluate the effect of ligand binding on the allosteric network (Fig. 3A).

**Figure 3.**
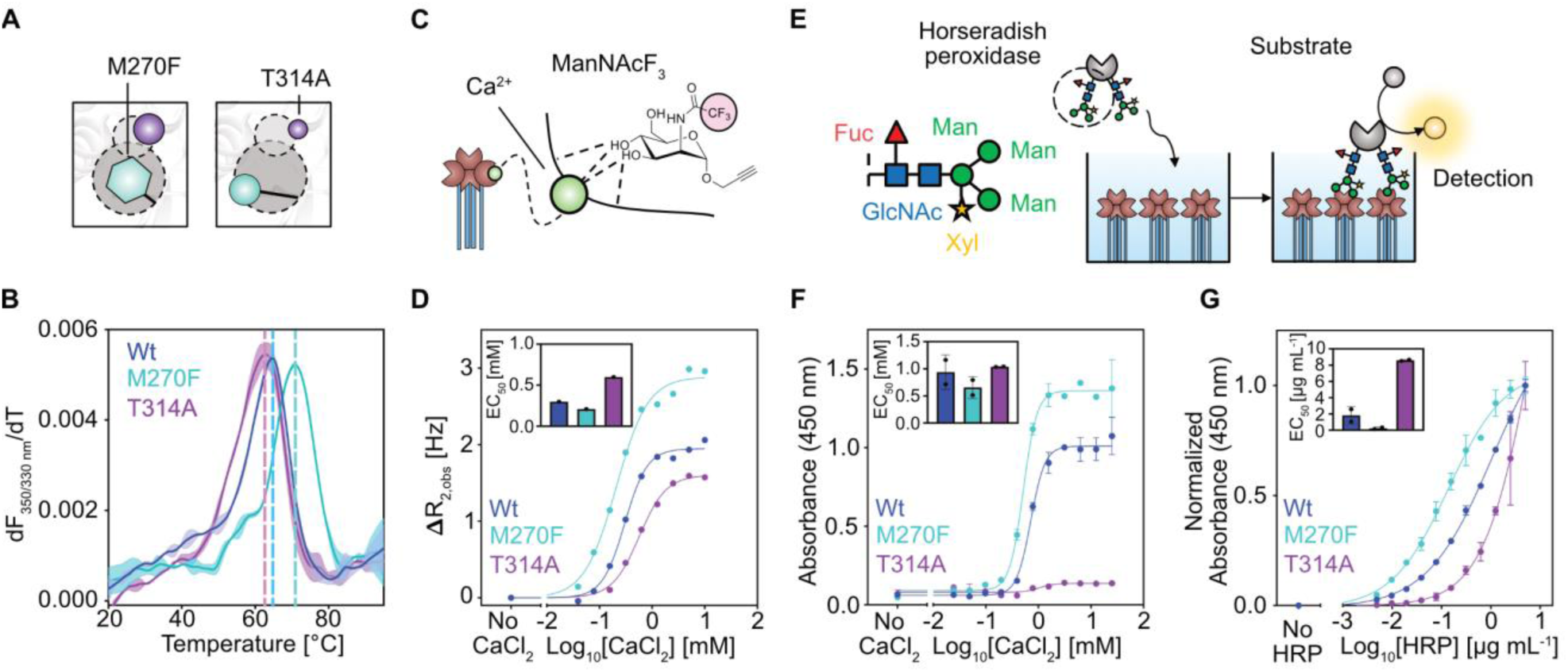
Mutations of network hubs T314 and M270 alter glycan binding properties of DC-SIGN. (**A**) The M270F mutant mimics a small molecule occupying the cryptic pocket. T314 lays in the upper cavity and was found to be a key network hub of the holo network. (**B**) The M270F mutant increases stability of the CRD. Melting curves of the ECD proteins in 25 mM HEPES, 150 mM NaCl, 10 mM CaCl2, pH 7.4. Inflection points (dotted lines) of the ratio of fluorescence recorded at 330 and 350 nm indicate the derived melting temperature T_m_. (**C**) Ca^2+^-dependent interaction of the fluorinated N-acetyl mannosamine analogue reporter (ManNAcF_3_) with the canonical carbohydrate binding site. Measurement of the binding to DC-SIGN ECD via T_2_-filtered ^19^F NMR enables indirect measurement of Ca^2+^ binding to the canonical carbohydrate binding site. (**D**) Binding of the reporter under varying CaCl_2_ concentrations to the ECD proteins reveals minor differences in Ca^2+^ affinity. (**E**) Glycans of the HRP enzyme bind to DC-SIGN ECD immobilized on a plate in a multivalent manner. Binding is evaluated using the peroxidase activity of HRP. (**F**) Binding of HRP at constant concentration to the ECD proteins at varying CaCl_2_ concentrations, reveal marginal change in Ca^2+^ affinity due to the mutations. (**G**) Binding of HRP at varying concentrations to the ECD proteins at 25 mM CaCl_2_, reveal significant impact of the mutations of glycan binding abilities of DC-SIGN. Means and standard deviations of EC_50_ values obtained from the HRP assay were calculated from two biological replicates each conducted in two technical replicates.

Considering their proposed importance in the Ca^2+^-dependent network, we hypothesized that the T314A and M270F mutations could affect folding, oligomerization and Ca²⁺ affinity as well as the associated glycan binding characteristics of DC-SIGN. Successful expression, refolding and purification of the ECDs *via* mannan affinity chromatography indicated the mutants to be correctly folded and in their active tetrameric state under Ca²⁺ saturating conditions. This was further confirmed by differential scanning fluorimetry (DSF) and dynamic light scattering (DLS) measurements indicating high thermal stability and tetramerization of the purified proteins (Figs. 3B and S17). Nevertheless, while the M270F mutant showed increased thermal stability (Tm = 70.4 ± 0.6 °C) compared to the wildtype (Tm = 64.5 ± 0.1 °C), the T314A mutation slightly decreased thermal stability (Tm = 63.0 ± 0.4°C). To study the effect of the mutations on Ca²⁺-dependent carbohydrate binding activity, we utilized our previously established ^19^F NMR reporter assay and measured binding of the fluorinated N-acetyl mannosamine analogue reporter (ManNAcF_3_) at a constant concentration under varying CaCl_2_ concentrations (*49, 57*). Importantly, as the binding mode of the reporter is dependent on Ca²⁺ coordination, this setup allows for measurement of an apparent affinity value for Ca²⁺ at the carbohydrate-coordinating Ca²⁺ site II by means of an EC_50_ value (Fig. 3C). Although Ca²⁺ affinity of DC-SIGN was not reported so far, the obtained EC_50_ for the wildtype protein (EC_50_ = 0.3 mM, hill slope = 2.0) was in agreement with previously reported affinities for the related CTL langerin (Fig. 3D) (*38*). Moreover, similar to other CTLs interacting with accessory Ca²⁺ ions in the long loop region, the fitted hill slope of >1 indicated positive cooperative effects for Ca²⁺ binding (*58, 59*). While this positive cooperativity was maintained in the mutants, the apparent Ca²⁺ affinity was slightly increased for DC-SIGN M270F (EC_50_ = 0.2 mM, hill slope = 1.4) and decreased for DC-SIGN T314A (EC_50_ = 0.6 mM, hill slope = 1.7). Moreover, these changes in affinity were accompanied by differences in the maximum R_2,obs_ values at saturating CaCl_2_ concentrations, with M270F showing higher R_2,obs_ than the wildtype and T314A showing lower R_2,obs_.

To orthogonally verify the changes in Ca²⁺-dependent carbohydrate binding, we titrated CaCl_2_ to the plate-immobilized ECDs in the presence of the tri-mannose-glycan carrying horseradish peroxidase (HRP) and detected binding using its peroxidase activity (Fig. 3E) (*60*). While the CaCl_2_ EC_50_ of the wildtype (EC_50_ = 1.0 ± 0.4 mM), the M270F (EC_50_ = 0.7 ± 0.2 mM) and the T314A mutant (EC50 = 1.2 ± 0.3 mM) remained similar to observations from the ^19^F NMR assay, T314A showed drastically reduced binding to HRP (Fig. 3F). As this could indicate differential glycan binding affinities, we titrated HRP at saturating CaCl_2_ concentration to the immobilized proteins and calculated its binding parameters. In line with the observed reporter and HRP binding in the CaCl_2_ titrations, the EC_50_ of T314A increased 4-fold (EC_50_ = 8.6 ± 0.1 µg mL^-1^) while the M270F mutant showed 10-fold reduced EC_50_ for HRP (EC_50_ = 0.2 ± 0.2 µg mL^-1^) compared to the wildtype (EC_50_ = 1.8 ± 1.0 µg mL^-1^) (Fig. 3G). Finally, we confirmed the glycan specificity of this interaction by competition experiments with mannose at constant HRP concentrations, revealing similar IC_50_s for all three proteins (Fig. S18).

Collectively, our binding assays demonstrated that mutations of the hub residues only marginally affect Ca²⁺ binding affinity but instead have a more significant effect on glycan binding. As we could exclude the reduced binding of T314A to HRP to originate from erroneous tetramerization of the T314A ECD from our DLS measurements, we hypothesized that T314 acts as a hub for regulating monovalent CRD-glycan interactions. Concomitantly, increased HRP binding upon occupation of the pocket, as mimicked by the M270F mutation, suggested the secondary site to activate carbohydrate binding, further supporting a model of an allosteric network modulating glycan binding in DC-SIGN.

### The allosteric network remodels the extended carbohydrate binding site

Our previous titrations showed only minor differences in mannose affinities between the T314A and M270F mutants and wildtype (Fig. S16). Direct comparison of the titration spectra further confirmed similar interaction modes, with nearly identical CSP trajectories for residues such as E347 and N349 of the EPN motif, which directly coordinate mannose at the canonical Ca²⁺-dependent binding site (Fig. 4A, C). This indicated that altered glycan-binding behavior in the mutants cannot be fully explained by changes at the canonical site. Instead, we observed that several residues within the extended carbohydrate binding site, including F313, E358, F359 and S360, exhibited pronounced CSPs and altered trajectories upon mannose titration, particularly in the T314A mutant (Fig. 4B, C). These residues, which are located in β-strands 3 and 4 and ⍺-helix 2, were also identified as hubs in our NMI analysis. On the one hand, this further confirmed allosteric coupling between the cryptic site, the upper cavity and the hubs identified in our MD simulations. On the other hand, we hypothesized that changes in binding to HRP, which carries core(1–3)fucosylated, xylosylated, trimannosyl oligosaccharides, could indicate the allosteric network to remodel the extended carbohydrate binding site, and therefore oligosaccharide binding (*61*).

**Figure 4.**
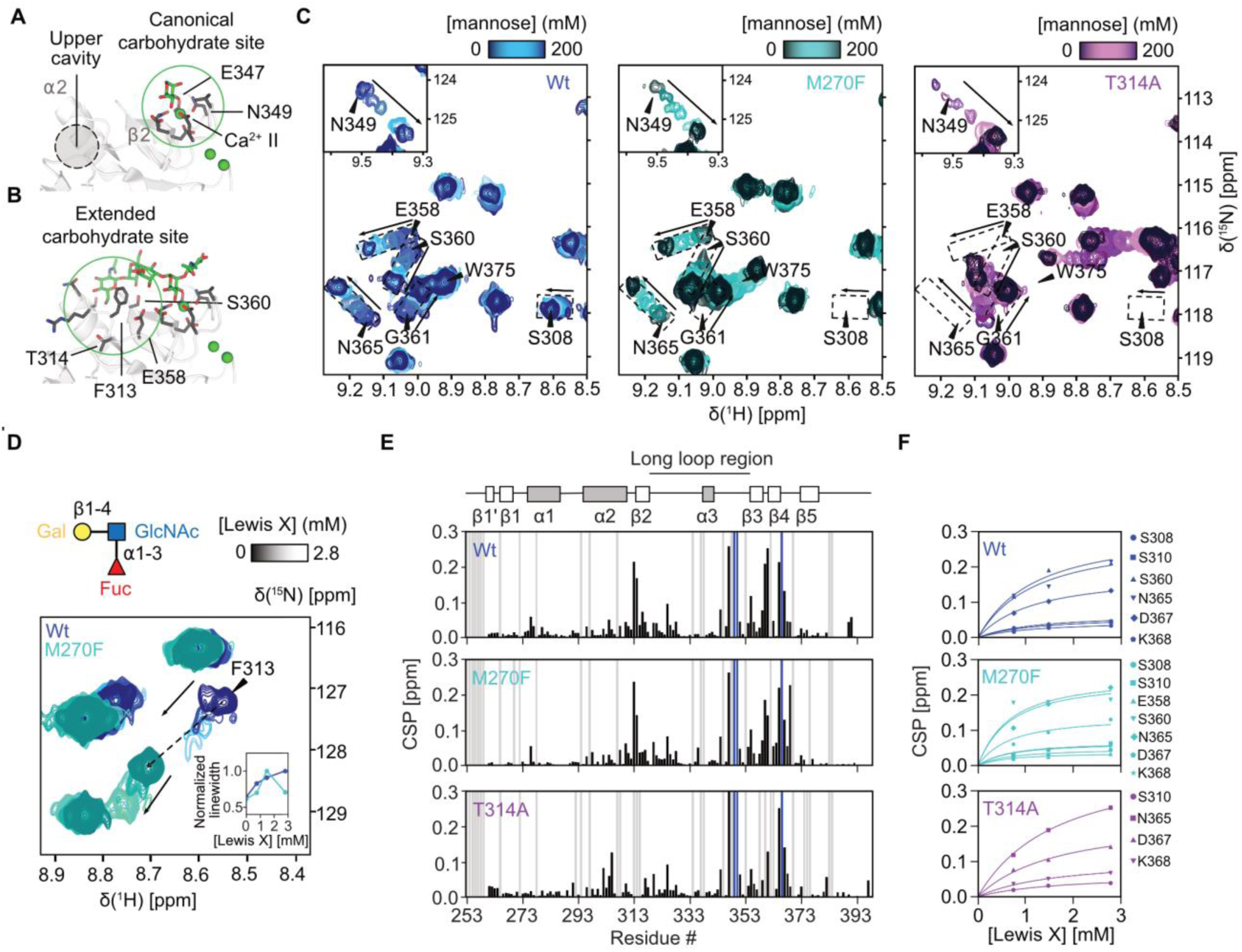
Mutations remodel the extended carbohydrate binding site. (**A**) Structural model of the canonical carbohydrate binding site formed by Ca^2+^ site II and E347 and N349 of the EPN motif. (PDB ID: 1K9I). Glycans apart from mannose monosaccharide were removed). (**B**) The extended carbohydrate binding site formed by β-strands 2-4, ⍺-helix 2 and the ⍺2-β2 connecting loop facilitates interactions with oligosaccharides such as GlcNAc_2_Man_3_ (PDB ID: 1K9I). (**C**) In line with similar mannose affinities, ¹H-¹⁵N HSQC NMR mannose titrations to the CRDs show only minor structural effect of the mutations on residues at the canonical carbohydrate site such as N349. In contrast, larger perturbations and changes in CSP trajectories in residues such as E358 and S360, suggest remodeling of the extended carbohydrate binding site. CSP trajectories of the wildtype are shown as boxes and projected onto the spectra of the mutants. (**D**) Comparison of ¹H-¹⁵N HSQC NMR shift trajectory of residue F313 upon titration of the Lewis X oligosaccharide to wildtype and M270F CRD. For both proteins F313 is in intermediate exchange but faster recovery of the linewidth indicates M270F to saturate faster than the wildtype protein. (**E**) CSP maps of Lewis X interacting with wildtype and the mutant CRDs. (**F**) Fitting of CSP trajectories reveals M270F (K_D_ = 0.6 ± 0.2) to increase affinity for Lewis X while T314A (K_D_ = 1.4 ± 0.1) has decreased affinity for Lewis X compared to the wildtype CRD (K_D_ = 1.1 ± 0.2).

To verify our hypothesis, we selected the Lewis X trisaccharide as it has been previously shown to interact with the extended carbohydrate site of DC-SIGN (Fig. 4D) (*62*). Using ¹H-¹⁵N HSQC NMR titrations with all three proteins, we compared CSP trajectories and dissociation constants. Addition of Lewis X to the wildtype induced large CSPs in similar residues as mannose (Figs. 4E, S19 and S20). However, several resonances of residues that were in fast exchange in mannose titrations displayed severe line broadening, indicative of intermediate exchange phenomena, suggesting higher affinity of Lewis X compared to mannose. In line with a previously proposed binding mode, this included residue F313 in the ⍺2-β2 loop pointing towards stronger involvement of the extended carbohydrate site in the Lewis X-DC-SIGN interaction (Fig. 4D). Although saturation was not reached at ∼3 mM Lewis X, estimating the dissociation constant from fast exchanging resonances yielded a K_D_ consistent with previously reported values (K_D_ = 1.1 ± 0.2) (Fig. 4F) (*62*). Strikingly, when compared to experiments with M270F, the F313 resonance in intermediate exchange saturated faster, as visible from changes in linewidth over ligand concentration (Fig. 4D). In line with this observation, the K_D_ was reduced two-fold (K_D_ = 0.6 ± 0.2) compared to the wildtype, indicating M270F to interact stronger with the trisaccharide. Conversely, we observed a lower K_D_ of the ligand for T314A (K_D_ = 1.4 ± 0.1) (Fig. 4F). Due to the significant change in chemical shift, we were unable to unambiguously transfer the resonance assignment for F313 from the wildtype to the T314A spectrum. Nevertheless, no additional resonances in intermediate exchange were observed upon addition of Lewis X, suggesting T314A to bind weaker to the oligosaccharide than the wildtype and M270F, further confirming observations from our HRP assay (Fig. S19). Moreover, changes in CSP trajectories of residues at the extended carbohydrate binding site and increased CSPs at residues in ⍺-helix 2 an β-strand 1’ hinted towards a different binding mode of the T314A-Lewis X complex (Figs. 4E and S20).

Taken together, comparative analysis of mannose and Lewis X binding to DC-SIGN wildtype and the mutants suggested that monosaccharide recognition at the canonical Ca²⁺-dependent site remains largely unaffected, while the allosteric network modulates oligosaccharide binding through structural rearrangements in the extended carbohydrate binding site. Finally, an increase in affinity of M270F and a decrease in affinity of T314A for Lewis X, likely by altering binding modes and exchange dynamics, provided further evidence for the cryptic pocket and the upper cavity to be functionally linked to the extended carbohydrate-binding site.

### Network hub mutations shift the Ca²⁺-dependent conformational equilibrium of ⍺-helix 2

Our binding studies demonstrated skewing of the interaction of DC-SIGN with oligosaccharides towards lower or higher affinity by inserting mutations in the network. If this observation results from perturbation of the same allosteric network, their effects should be reflected in a concerted conformational response across structural elements involved in allosteric modulation (*63*). To test this, we superposed the ¹H-¹⁵N HSQC NMR spectra of wildtype, M270F, and T314A and compared direction and magnitude of chemical shift changes. Several resonances displayed linear displacement alongside a unifying chemical shift vector (Fig. S21). As chemical shifts represent the population-weighted averages of conformational equilibria in the fast exchange regime, this linearity suggested exchange between two conformational states (*55*). The linear chemical shift correlation was most pronounced for residues T261, F262, F263, Q264 around the cryptic site in β-strand 1’, Q300 and F302 in ⍺-helix 2 and R312 in the ⍺2-β2-connecting loop (Fig. 5A). Structurally, F263 and F302 are involved in an orbital π-stacking interaction, anchoring β-strand 1’ onto the rest of CRD structure through ⍺-helix 2. Similar π-stacking interactions have been implicated in the oligomerization behaviour of the related C-type lectin langerin (*64*). Yet, DC-SIGN and langerin oligomers display distinct topology and relative orientation of CRDs and we observed both mutants to form tetramers similar to the wildtype (*28, 65*). While it is possible that this interaction impacts the relative spatial arrangement CRDs without affecting tetramerization via the neck domain, changes in affinity of both ECD and CRD suggested intradomain allosteric control (*53*). In support of this, close packing of the C-terminal end of ⍺-helix 2 and the ⍺2-β2 connecting loop formed by N311, R312, F313 and T314 towards β-strands 3 and 4 has been previously described to provide a continuous surface for recognition of oligosaccharides in the extended carbohydrate binding site (*25*). While this could point to a mechanism in which oligosaccharide binding could be modulated by the relative positioning of ⍺-helix 2 towards β-strands 3 and 4, several residues at the C-terminal end of ⍺-helix 2 towards the upper cavity deviated from co-linearity indicating the presence of at least one additional state in that region (Figs. 5A and S21) (*66*). Moreover, residues that exhibited linear displacement showed only minor shifts in the Lewis X titrations with the wildtype, suggesting that their structural transitions are not primarily driven by oligosaccharide binding (Fig. 4E).

**Figure 5.**
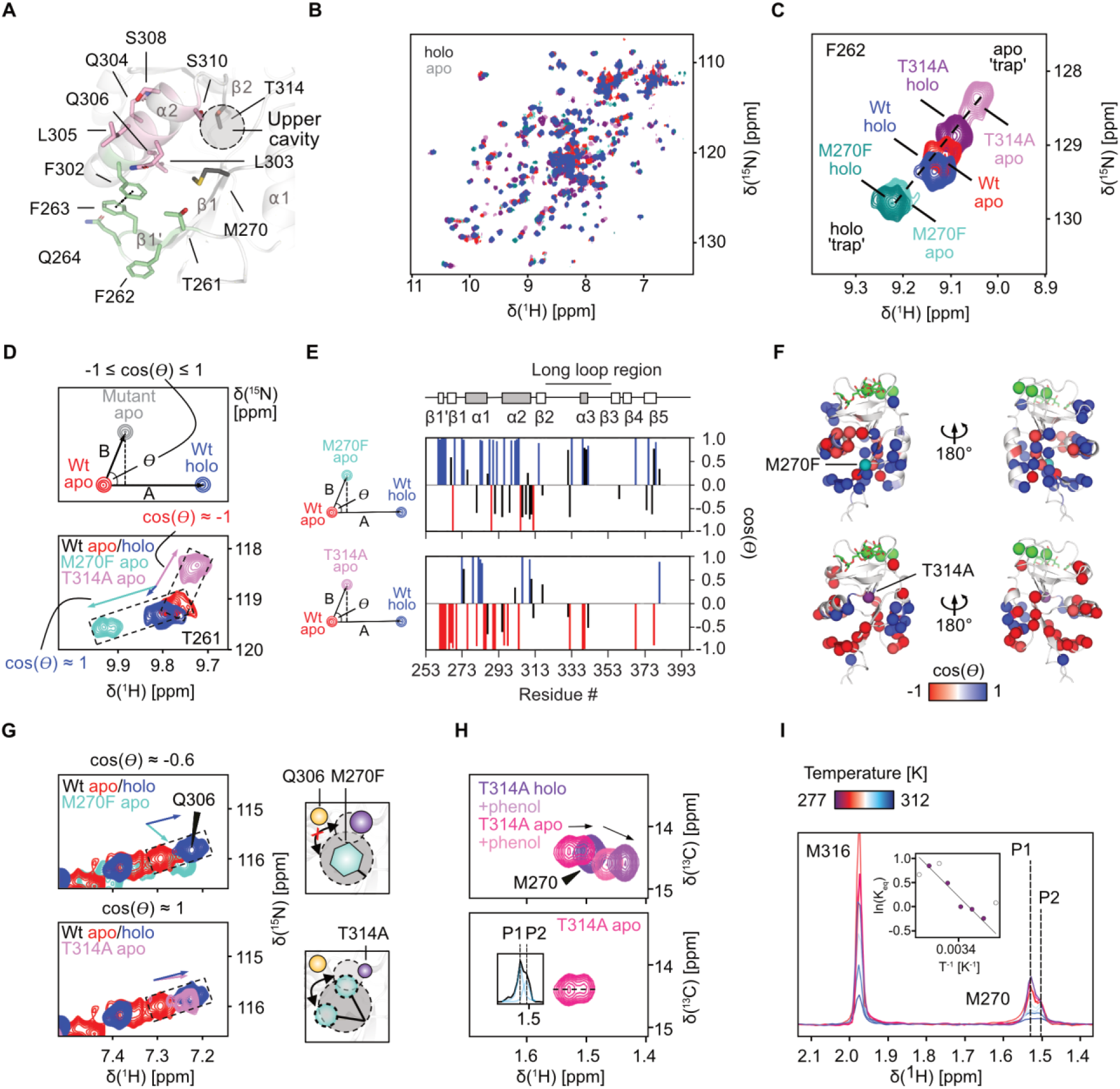
Mutations shift the conformational equilibrium of DC-SIGN by altering the open-closed equilibrium of the cryptic pocket. (**A**) Residues of the secondary site and ⍺-helix 2 that show co-linear displacement in ¹H-¹⁵N HSQC NMR spectra along a unifying shift vector upon T314A and M270F mutation (light green sticks). Towards the C-terminal end of ⍺-helix 2 and the upper cavity residues show non-linear behavior (red sticks). Spectra are shown in Fig. S21. Residue F263 and F302 engage in an orbital π-stacking interaction (black dotted line), anchoring β-strand 1’ through ⍺-helix 2 onto the rest of the CRD. (**B**) Overlay of apo and holo ¹H-¹⁵N HSQC NMR spectra of the wildtype and the mutant CRDs. (**C**) Example of residue β-strand 1’ showing continued co-linearity in chemical shift in apo and holo spectra of the wildtype and the mutants. The T314A mutation shifts the resonance beyond the apo state even in the presence of Ca²⁺ (apo ‘trap’), while M270F shifts the resonance beyond the holo state (holo ‘trap’). (**D**) Vector scheme of CHESPA to determine the angle θ between ¹H-¹⁵N HSQC NMR chemical shift vectors of apo to holo wildtype and apo wildtype to apo mutant. A cos(θ) ≈ 1 indicates conformational similarity to holo wildtype, while cos(θ) ≈ -1 suggests similarity to the apo wildtype conformation. The resonance of residue T261 is shown as an example. (**E** and **F**) Comparison of all cos(θ) values suggests M270F (top) to shift the CRD towards the holo state, while T314A induces a shift towards the apo state. Residues of the upper cavity deviate from this trend showing a shift towards the holo conformation for T314A and towards the apo conformation for M270F. (**G**) Comparison of the Q306 chemical shift of the apo mutants and apo and holo wildtype suggest T314A to induce an holo-like state in this residue (cos(θ) ≈ 1), while the Q306 resonance of M270F deviates from co-linearity (cos(θ) ≈ -0.6). (**H**) ¹H-^13^C HSQC NMR spectra of apo and holo ^13^C Met-labeled DC-SIGN CRD T314A interacting with phenol. The CSPs induced by Ca²⁺ do not follow a unifying shift vector as observed for the wildtype (Fig. 2C). The apo spectrum shows a split resonance for M270, indicating two slowly interchanging states P1 and P2. The later resembles the holo state of M270. The inset shows the line fitting of the extracted ¹H dimension of the apo spectrum, indicating ∼35% of the protein to be in state P2. (**I**) Line fitting of the extracted ¹H dimension of the apo spectrum at varying temperatures shows a shift towards P1. The inset shows fitting of the populations over temperature in a Van’t Hoff plot, suggesting an endothermic transition (ΔH = +32 kJ mol^-1^) and a positive entropy change (ΔS = +112 J mol-1 K^-1^). Uncolored data points were not used for fitting. ¹H-^13^C HSQC NMR spectra of the resonance at different temperatures are shown in Fig. S22.

Given that our MD simulations and the Ca²⁺ titration indicated that structural changes upon Ca²⁺ binding could potentially propagate to the upper cavity and the cryptic pocket via ⍺-helix 2, we revisited this interaction and evaluated the relationship to our mutants by comparing apo and holo ¹H-¹⁵N HSQC NMR spectra of all three proteins (Fig. 5B). Intriguingly, we observed continuation of the linear shift trajectories in several residues, such as F262, F263, Q264 and F302 demonstrating a concerted structural response to the M270F and T314A mutations and Ca²⁺ binding. In line with our MD simulations, this suggested that a Ca²⁺-responsive network is at play that can be perturbed by mutating hub residues. Along trajectories of resonance of residues at the interface of β-strand 1’, α-helix 2, and the cryptic site, the apo M270F resonances shifted beyond the respective holo wildtype resonances, suggesting the mutant to stabilize a holo-like state, while the T314A mutation appeared to stabilize an apo-like state. As neither M270F, nor T314A recovered the wildtype holo state upon removal or addition of Ca²⁺ in these resonances, respectively, we speculated that the effect induced by the mutations could ‘trap’ ⍺-helix 2 in a either holo or apo-like conformation independent of Ca²⁺ (Fig. 5C).

### The T314A mutation shifts the open-closed equilibrium of the cryptic site towards the open state

Comparative chemical shift analysis revealed a concerted linear response of the CRD to Ca²⁺ binding and the mutations in β-strand 1’, α-helix 2, and the cryptic site suggesting a two-state equilibrium in this region. However, deviations from this linearity in several residues indicate partial decoupling towards the C-terminal end of ⍺-helix 2 and the upper cavity that could hold additional information on the origin of the allosteric effect of the cryptic site.

To evaluate this systematically we compared the conformational similarity of holo and apo wildtype and the apo mutant spectra using chemical shift projection analysis (CHESPA) of the ¹H-¹⁵N HSQC NMR spectra (*67*). CHESPA involves a simple comparative vectorial analysis of CSPs resulting from mutation-induced perturbations and a reference perturbation caused by an orthosteric ligand or co-factor. Based on the cosine of the angle θ between the shift vectors, the degree of conformational similarity to the activated protein bound to an orthosteric ligand or co-factor can be quantified, revealing whether distinct structural states project onto a common binding-dependent equilibrium (Fig. 5D). Although intermediate exchange phenomena in apo ¹H-¹⁵N HSQC NMR spectra, especially of residues at the Ca²⁺-dependent canonical and the extended binding site, is a key limitation of this approach, we reasoned that peripheral effects could allow us to evaluate the relationship between the mutants and Ca²⁺ binding. As expected, resonances of residues in β-strand 1’, α-helix 2, and the cryptic site approached cos(θ) ≈ 1 for M270F but showed cos(θ) ≈ -1 for T314A (Fig. 5E, F). However, these effects did not apply to the C-terminal end of ⍺-helix 2 and the upper cavity, as residue Q306 showed cos(θ) ≈ 1 for apo T314A and a non-linear arrangement with cos(θ) ≈ -0.6 for M270F (Fig. 5G). Since Q306 presumably stabilizes the open state of the cryptic pocket by forming a hydrogen bond to the thioether sidechain of M270, we hypothesized that this observation could reflect the inability of the M270F mutant to open the cryptic pocket by rotation of residue 270 into the upper cavity (Fig. S4). Contrastingly, as we have observed the M270 sidechain to resemble the holo state upon phenol binding to apo DC-SIGN wildtype in our ¹H-^13^C HSQC NMR experiments, we reasoned that the holo-like state of Q306 in T314A could indicate a shift in the open-closed equilibrium towards the open species, therefore exposing the hydrophobic cryptic pocket even in the absence of a ligand (Fig. 5G).

As the open-closed conformational equilibrium, and with it the chemical shift of Q306, should be coupled the M270 sidechain dynamics, we recorded ^1^H-^13^C HSQC NMR spectra of apo and holo ^13^C Met-labeled T314A in the presence and absence of phenol and compared them to experiments with the wildtype. In line with our hypothesis of a preferred open state upon mutation, we observed a shift of the M270 resonance in holo T314A beyond the resonance of the wildtype bound to phenol. In the presence of phenol, the T314A trajectory is continued with a CSP of higher magnitude, suggesting that T314A preserves the ability to interact with ligands at the cryptic site and that this interaction might be of higher affinity (Fig. 5H). Intriguingly, removing Ca²⁺ from T314A, revealed a split in the M270 resonance, with a higher populated state that resembled the wildtype bound to phenol and a second state overlapping with the holo T314A M270 resonance. This aligns with our CHESPA data, suggesting that even in the absence of Ca²⁺ M270 adopts a conformation resembling that of the holo state. Moreover, the presence of two distinct peaks for M270 in the absence of Ca²⁺ suggests that these conformations interconvert in the slow exchange regime on the NMR timescale. Analysis of the temperature-dependent population shifts revealed that both states remain in near-equilibrium up to ∼298K, after which the apo state becomes increasingly populated, suggesting an entropically driven transition (Figs. 5I and S22). Van’t Hoff analysis confirmed that this shift is thermodynamically controlled, with an endothermic transition (ΔH = +32 kJ mol^-1^) and a positive entropy change (ΔS = +112 J mol^-1^ K^-1^), indicating that while the holo state is enthalpically stabilized at lower temperatures, the release of Ca²⁺ leads to an increase in conformational freedom of the cryptic pocket.

Together with the CHESPA results and the slow-exchange behavior, the thermodynamic profile strongly suggested that the Ca²⁺-dependent conformational state of M270 in apo T314A is already biased toward an open-pocket conformation, partially decoupling the phenol bound from the Ca²⁺ bound conformation. The decoupling of this conformation from phenol binding was further supported by the non-linearity of the CSPs when comparing spectra of apo and holo T314A with and without phenol. Unlike in the wildtype, where Ca²⁺ and phenol binding shift along a unifying vector, a loss in correlation is observed in the CSP pattern of T314A (Figs. 5H and S9C). Instead, phenol binding perturbs an already open conformation, reinforcing the idea that T314A shifts the energetic landscape of the cryptic pocket to favor its open state.

### Retrospective analysis of X-ray structures suggests a stabilizing effect of ligand binding at the cryptic pocket

Our NMR data strongly suggested the mutants T314A and M270F to favor an open state or mimic an occupied state devoid of opening of the cryptic pocket, respectively. This also implies that the concerted chemical shift response to Ca²⁺ and the mutations in β-strand 1’ and α-helix 2 observed in ^1^H-¹⁵N HSQC NMR could directly report on the populations of the exposed and bound state of the cryptic pocket (*68*). This would indicate that binding to the pocket could shift the structure to a rather Ca²⁺ bound state, which would, according to our model, lead to a stabilization of the extended carbohydrate binding site. In turn, exposure of the hydrophobic pocket to the solvent as seen in T314A, would destabilize the extended carbohydrate binding site. Nevertheless, in an actual ligand binding event to the wildtype, a mixture of the T314A and M270F conformations is expected, with M270 rotating into the upper cavity and the ligand occupying the cryptic pocket.

To date, no high affinity ligand or co-crystal structure of a ligand-occupied cryptic site is available. Yet, we reasoned that changes in stability and conformation should be reflected in the comparison of the X-ray structures of wildtype CRD in the open and closed conformation. Comparison of the normalized B factors in wildtype CRD X-ray structures in the open (PDB ID: 1SL5) and closed (PDB ID: 1SL4) conformations revealed that pocket opening alters structural flexibility (Fig. 6A) (*52*). Normalized B factors decreased at the N-terminal end of ⍺-helix 2 but increased at its C-terminal end, forming the upper cavity. While the latter could result from the M270 sidechain competing water from the upper cavity, decreased normalized B factors of residues in ⍺-helix 2 suggest increased stability upon pocket opening and binding of the tryptophan sidechain at the cryptic pocket (Figs. 6B and S4). Similarly, decreased B factors in the loop and β-strands 3 and 4 suggest stabilization at the extended carbohydrate binding site. Finally, in support of a stabilizing effect of closing the cryptic pocket, we found that phenylalanine seems to be evolutionary favored across human C-type lectins at the position equivalent to M270 in DC-SIGN (Fig. S23).

**Figure 6.**
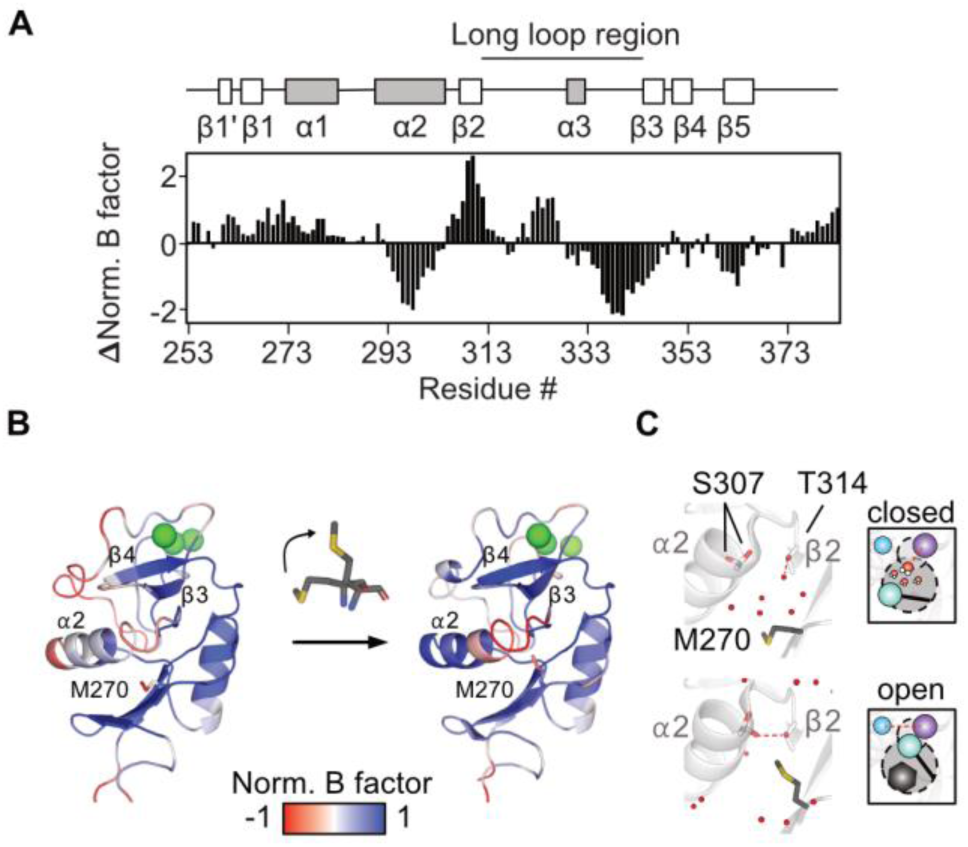
Ligand binding at the cryptic pocket stabilizes the CRD. (**A**) Difference of normalized B factors extracted from C⍺ atoms in the X-ray crystallographic structures of DC-SIGN CRD in the closed (PDB ID: 1SL4) and in the open state (PDB ID: 1SL5). Lower values indicate residues with higher B factor in the closed form and *vice versa*. (**B**) Normalized B factors extracted from all atoms of the same structures and mapped back onto the structures display changes in stability around ⍺-helix 2 and β-strands 3 and 4. (**C**) Close up of the upper cavity in the closed (top, PDB ID: 1SL4) and the open form (bottom, PDB ID: 1SL5) displaying changes in the hydrogen bond pattern upon opening of the cryptic pocket.

Overall, changes in stability were not accompanied by major backbone rearrangements. However, in accordance with our phenol-MD simulations, water competition in the upper cavity altered the hydrogen bonding pattern of T314 (Fig. S1B). In the closed state, T314 forms a hydrogen bond to a water molecule, while rotation of M270 in the open state removes water, enabling a T314–S307 hydrogen bond in ⍺2 (Fig. 6C). Despite the flexibility of the S307 sidechain, this may stabilize interactions between ⍺-helix 2 and β-strand 2 and influences the adjacent ⍺2–β2 loop bearing residue F313 involved in oligosaccharide binding. Our apo-holo MD simulations showed this bond to occur exclusively in the holo state while being absent in the apo state (Table S1). Consequently, the T314A mutant likely reduces water occupancy in the upper cavity, allowing M270 rotation and favoring the open conformation even in the absence of a ligand. Moreover, it precludes the T314-S307 hydrogen bond, which may also destabilize the holo state, trapping ⍺-helix 2 in its apo conformation. Although no apo X-ray structure exists, our MD simulations also suggested a hydrogen bond between Q300 (⍺2) and W364 (β4, WND motif) to be weakened in the apo state. The WND motif links Ca²⁺ coordination at the glycan-binding site to the hydrophobic core and, as our data suggest, also to ⍺-helix 2 (*26*).

## Discussion

DC-SIGN-glycan interactions are critical for pathogen recognition, immune modulation, and cellular adhesion (*4, 5, 31, 33, 69, 70*). Uncovering how these interactions are regulated at a molecular level offers critical insight into how glycan recognition by the DC-SIGN CRD translates into downstream cellular response, thereby advancing our understanding infectious diseases and immune disorders, eventually informing DC-SIGN-targeted drug discovery. Past work has identified a secondary site that was proposed to allosterically activate carbohydrate binding in DC-SIGN through interactions with a bivalent glycomimetic ligand, but the underlying mechanism remained elusive (*49*). Here, we demonstrate that binding to this site underlies cryptic pocket opening, allosterically activating glycan binding *via* the extended carbohydrate binding site of DC-SIGN.

We propose a model wherein the occupation state of the cryptic pocket modulates glycan binding affinity through conformational changes in ⍺-helix 2, adopting either a holo-like or apo-like state depending on pocket occupancy. Central to this mechanism is M270, a gatekeeper residue whose sidechain rotation into an upper cavity enables cryptic pocket opening, exposing a hydrophobic cleft. We identified the M270F and T314A mutations as proxies for the occupied and unoccupied states of the cryptic pocket, increasing or decreasing glycan affinity, respectively. These phenotypes are linked to a concerted structural response, skewing the Ca²⁺-dependent conformational equilibrium of DC-SIGN, specifically affecting ⍺-helix 2. While T314A traps this region in an apo-like state, M270F stabilizes a conformation resembling the holo-state of the wildtype, even in the absence of Ca²⁺. The helix therefore behaves as a two-state hinge coupling pocket occupancy to glycan recognition.

While interactions with the Ca²⁺-coordinated EPN motif remain largely unchanged in the mutants, our NMR studies, MD simulations, and glycan-binding assays point to the extended carbohydrate binding site as the major site of allosteric regulation. Residues at the interface of α-helix 2 and β-strands 2–4, including F313, S360, and E358, show pronounced chemical shift perturbations upon mutation and were identified as high-connectivity nodes in the network. Previous structural studies have shown that the same residues contribute to glycan specificity and affinity by forming secondary contacts that enable DC-SIGN to accommodate oligosaccharides in distinct geometries (*25, 27*). In contrast, interactions at the canonical site are less variable, and, independent of the glycan involved, uniformly interact with the same hydrogen bond acceptors and donors at the canonical carbohydrate binding site via their 3-OH and 4-OH groups (*71*). These observations suggest that the allosteric mechanism we describe modulates glycan affinity and selectivity by priming, potentially reshaping the extended binding site.

We found Ca²⁺ complexation to assume a structural role beyond direct interactions with monosaccharides. Both our MD simulations and NMR studies showed apo DC-SIGN to sample a broader conformational ensemble, similar to what has been described for the related C-type lectins DC-SIGNR and langerin (*38, 72*). Binding to Ca²⁺ leads to a reduced rate of interconversion between conformations and to higher connectivity, especially affecting residues of the extended site and ⍺-helix 2 but also the open-closed conformation of the cryptic pocket. Accordingly, co-factor complexation potentially has a similar effect on DC-SIGN as occupation of the cryptic site - that is priming the extended carbohydrate binding site for interactions with oligosaccharides. We hypothesize, that this effect is mediated by a network of interactions, including a W364-Q300 hydrogen bond that links the Ca²⁺ cage via the WND motif to α-helix 2, an orbital π-π interaction between F263 and F302 anchoring β-strand 1’ and a T314-S307 hydrogen bond that likely stabilizes the holo-like state (*26*). Supporting this mechanism, pocket occupation results in reduced flexibility in α-helix 2 and β-strands 3 and 4 upon pocket occupation and increased stability as suggested by melting temperatures and evolutionary preference of a phenylalanine in position of M270. While reduced flexibility of the carbohydrate binding site as affinity driving factor in C-type lectins has not been investigated so far, studies on other glycan binding proteins, for instance galectins, have shown that a preorganized carbohydrate recognition domain and conformationally-restricted glycan ligands can substantially diminish the entropic penalty associated with binding of conformationally flexible glycans, thereby enhancing overall affinity and shaping selectivity (*73, 74*).

While the dominant two-state equilibrium is centred around α-helix 2, several observations also pointed towards extensive dynamics connecting the canonical carbohydrate binding site, the extended site and the Ca²⁺ sites. First, residues such as E358, S360, and F313 show large mutation-induced CSPs that did not follow linear chemical shift behaviour, suggesting the presence of intermediate states beyond a simple two-state model (*66*). Second, these same residues are also perturbed upon monosaccharide binding, despite being spatially distant from the canonical Ca²⁺-coordinated site, indicating that the canonical and extended sites are also structurally linked (*12*). Third, previous studies on the closely related C-type lectin DC-SIGNR have shown interdependence of Ca²⁺ binding at the accessory Ca²⁺ sites and glycan binding (*72, 75*). Therefore, subtle changes in Ca²⁺ affinity of DC-SIGN in our binding assays might also be rooted in our setup only indirectly measuring Ca²⁺ affinity *via* glycan binding. Finally, as we observed positive cooperativity between the Ca²⁺ sites of the CRD, mirroring observations for other C-type lectins such as ASGPR and MGL, several additional conformations of the long loop region and adjacent sites are possible (*58, 59*).

Taken together, our observations draw a complex picture of hierarchical allosteric modulation of DC-SIGN with both local and global dynamics differentially affecting the outcome of glycan binding. Previous studies on DC-SIGN have described a relationship between glycan binding dependent effects on both endocytosis and signaling (*3, 31, 33, 34, 76*). Thus, it is tempting to speculate whether the allosteric mechanism described here also translates into broader receptor-level response under physiological conditions as suggested for other C-type lectins, such as MGL and CLEC5A (*35, 36*).

Although the currently available data did not allow us to clearly separate the different effects and their contributions, the integration of an allosteric cryptic site into the network suggests that this complex interplay can be modulated by drug-like ligands. In this context, our results also highlight the potential of combining data from simulations with subsequent experimental validation by NMR to identify hidden cryptic sites in proteins with low-druggability. Cryptic pockets have emerged as useful entry points for ligand discovery targeting proteins with primary sites of low druggability (*77*). While secondary sites and in some cases allosteric mechanisms have been described for other C-type lectins, the present study is, to the best of our knowledge, the first description of a cryptic pocket and its functional effect identified in a glycan binding C-type lectin (*41, 42, 44, 46, 49, 78*). By enabling selective tuning of extended-site engagement without disrupting Ca²⁺ coordination or glycan specificity at the canonical site, targeting the here identified site could enable fine-tuning of cell-cell and cell-pathogen interactions, eventually tuning the immunological response to specific glycans. As other glycan-binding C-type lectins suffer from low druggability similar to DC-SIGN, we envision our approach to equally stimulate drug discovery campaigns targeting other pharmaceutically relevant members of this protein family (*40*).

## Materials and Methods

### Protein Expression and Purification

#### General remarks

Unless stated otherwise, all chemicals, growth media and enzymes used for protein expression and purification were purchased from Sigma Aldrich or Carl Roth. Codon-optimized genes for the bacterial expression of wildtype and mutant DC-SIGN CRD and ECD were purchased from GenScript.

#### DC-SIGN carbohydrate recognition domain

¹⁵N-labelled DC-SIGN CRD wildtype and mutants were produced as previously described (*49*). In brief, the protein was expressed insolubly in BL21 (DE3) *E. coli* transformed with a pET28a vector encoding amino acids 253–404 of DC-SIGN and a *N*-terminal His-tag. Bacteria grown in M9 minimal medium supplemented with 35 mg L^-1^ kanamycin and 0.5 g L^-1^ ¹⁵NH_4_Cl were induced at OD_600_ of 0.9 using IPTG, lysed and inclusion bodies were harvested by centrifugation. Following solubilization, the protein was refolded overnight *via* rapid dilution. Next, the protein was dialyzed against 50 mM Tris-HCl, 150 mM NaCl, 10 mM CaCl_2_ (pH 7.8) and purified *via* Ni^2+^ NTA affinity chromatography. Purified protein was pooled and dialyzed against 20 mM MES, 40 mM NaCl, 10 mM CaCl_2_ (pH 6.0), concentrated using a centrifugal spin filter, snap frozen and then stored at −80°C.

^13^C Met-labeled DC-SIGN CRD wildtype, M270F and T314A were produced following the same protocol as described for ^15^N-labeled DC-SIGN CRD, with the exception that 0.1 g L^-1^ ^13^C ε-methyl methionine was added to the M9 minimal medium at OD_600_ of 0.45 to avoid scrambling of the isotope.

#### DC-SIGN extracellular domain

DC-SIGN ECD wildtype, M270F and T314A were produced as previously described (*49*). In brief, the protein was expressed insolubly in BL21 (DE3) *E. coli* transformed with a pET30b vector encoding amino acids 64–404 of DC-SIGN. Bacteria were grown in Luria-Bertani (LB) medium with 35 mg L^-1^ kanamycin and expression was induced with IPTG at OD_600_ of 0.9. Following lysis, inclusion bodies were harvested by centrifugation and solubilized. The protein was refolded via rapid dilution and dialyzed against 25 mM Tris-HCl, 150 mM NaCl, 25 mM CaCl_2_, pH 7.8, for subsequent purification via mannan agarose affinity chromatography. Purified protein was dialyzed against 25 mM HEPES, 150 mM NaCl, 10 mM CaCl_2_, pH 7.4, concentrated using a centrifugal spin filter, snap frozen and then stored at -80°C.

^13^C ε-methyl methionine-labeled DC-SIGN ECD was expressed following the same protocol as described for DC-SIGN CRD by adding 0.1 g L^-1^ ^13^C ε-methyl methionine to the M9 minimal medium at OD_600_ of 0.45. Following expression, the protein was purified as described for the unlabeled ECD.

The tetrameric state of the purified proteins was confirmed using DLS measurements at 0.1 mg mL^-1^ protein concentration on a Zetasizer Advance Pro (Malvern Panalytical). Melting temperatures T_m_ were measured at 0.1 mg mL^-1^ protein concentration in thermal shift assays on a NanoTemper Prometheus (NanoTemper).

### Plate-based horseradish peroxidase assay

#### General remarks

Unless stated otherwise, all reagents and buffers were obtained from Sigma Aldrich or Carl Roth. Measurements were done on a Enspire Multimode Plate Reader (PerkinElmer). GraphPad Prism was used for all data processing and analysis.

#### CaCl_2_ titration

100 µg mL^-1^DC-SIGN ECD were immobilized in immobilization buffer (25 mM HEPES, 150 mM NaCl, 25 mM CaCl_2_, pH 7.4) overnight at 4°C on transparent Nunc Maxisorp 96-well plates (Thermo Fisher). Protein solution was removed, and plates were washed three times with immobilization buffer and then blocked for 2 hours at 4°C with blocking buffer (25 mM HEPES, 150 mM NaCl, 25 mM CaCl_2_, 2% BSA, 0.01% Tween-20). Plates were washed twice with 25 mM HEPES, 150 mM NaCl, 5 mM EDTA, pH 7.4 and twice with 25 mM HEPES, 150 mM NaCl, pH 7.4 to remove residual Ca²⁺. To titrate Ca²⁺, CaCl_2_ in 25 mM HEPES, 150 mM NaCl, pH 7.4 was serially diluted on the plate at a constant HRP concentration of 1 µg mL^-1^ and then incubated at RT for 2 hours. Following incubation, plates were washed three times with 25 mM HEPES, 150 mM NaCl, pH 7.4 and bound HRP was detected using a TMB substrate kit (Thermo Fisher) and 0.18 M H_2_SO_4_ according to manufacturer’s instructions. Absorption was measured at 450 nm.

#### HRP titration

Immobilization and blocking were done as described above. After blocking, plates were washed three times with immobilization buffer. HRP in immobilization buffer was serially diluted on the plate at a constant concentration CaCl_2_ concentration of 25 mM and then incubated at RT for 2 hours. Following incubation, plates were washed three times with immobilization buffer and HRP was detected as described above.

#### Mannose titration

Immobilization and blocking were done as described above. After blocking, plates were washed three times with immobilization buffer. Mannose in immobilization buffer containing 1 µg mL^-1^ HRP was serially diluted on the plate and then incubated at RT for 2 hours. Following incubation, plates were washed three times with immobilization buffer and HRP was detected as described above.

### NMR spectroscopy

#### General remarks

¹H-^13^C HSQC NMR, ¹H-¹⁵N HSQC NMR, ¹H-¹⁵N SOFAST-HMQC NMR and ^19^F NMR measurements were conducted on an Ultra Shield 500 MHz spectrometer (Bruker) equipped with a TCI H/F-C-N Prodigy probe. If not state otherwise all ¹H-^13^C HSQC NMR, ¹H-¹⁵N HSQC NMR and ¹H-¹⁵N SOFAST-HMQC NMR spectra were collected at 298 K. ¹H-^13^C TROSY NMR measurements were conducted on an Ascend 700 MHz spectrometer (Bruker) equipped with a TCI H/F-C-N Helium CryoProbe at 298 K. All samples were measured at sample volumes of 160 µL in 3 mm NMR tubes (Bruker). Unless stated otherwise, all reagents and buffers were obtained from Sigma Aldrich or Carl Roth. Lewis X trisaccharide was obtained from Biosynth. Spectra were processed in TopSpin 4.2.0 and data analysis was performed using CCPN Analysis 3.2.0 for 2D spectra and in MestreNova for 1D spectra (*79*). Further analysis was done using GraphPad Prism and in-house python scripts.

#### ¹H-^13^C HSQC NMR

^1^H-^13^C HSQC NMR spectra were collected with 512 increments in the carbon and 8 scans per increment and 2048 points in the direct dimension. The relaxation delay d1 was set to 1.5 s. The W5 WATERGATE pulse sequence was used for solvent suppression (*80*). Experiments were performed at a CRD concentration of 200 μM in 20 mM MES, 40 mM NaCl, 10 mM CaCl_2_ (pH 6.0) supplemented with 10% D_2_O and, if applicable, 20 mM phenol or 20 mM mannose. For spectra of apo DC-SIGN CRD and CaCl_2_ titrations the protein was dialyzed against 20 mM MES, 40 mM NaCl, 1 mM EDTA (pH 6.0) twice, and then against 20 mM MES, 40 mM NaCl (pH 6.0) twice prior to sample preparation and measurements. Reference spectra containing only buffer were used to evaluate potential scrambling of the ^13^C ε-methyl methionine (Fig. S3).

The temperature-dependent population shift of the M270 resonance was analyzed by extracting populations P1 and P2 by applying a general Lorentzian line fitting to the extracted ¹H projections of ^1^H-^13^C HSQC NMR spectra recorded at different temperatures. Thermodynamic parameters were obtained from fitting a Van’t Hoff plot according to **Equation 1**.

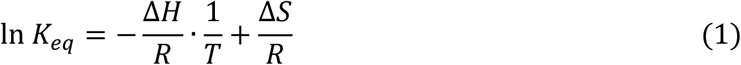

With ln K_eq_ as the natural logarithm of the equilibrium constant of the populations (P1/P2), ΔH the enthalpy change (in J mol^-1^), ΔS the entropy change (in J mol^-1^ K^-1^), R the gas constant (8.314 J mol^-1^ K^-1^) and T the absolute temperature in K.

#### ¹H-^13^C TROSY NMR

¹H-^13^C TROSY NMR spectra were acquired at a sample temperature of 298 K with 128 increments in the carbon and 16 scans per increment and 2048 points in the direct dimension. The relaxation delay d1 was set to 0.45 s. The W5 WATERGATE pulse sequence was used for solvent suppression (*80*). Experiments were performed at a ECD concentration of 300 μM in 25 mM HEPES, 150 mM NaCl, 10 mM CaCl_2_ (pH 7.4) supplemented with 10% D_2_O and, if applicable, 20 mM phenol.

#### ¹H-¹⁵N HSQC NMR

^1^H-^15^N HSQC NMR spectra were collected with 256 increments in the nitrogen and 24 scans per increment and 2048 points in the direct dimension. The relaxation delay d1 was set to 1.0 s. The W5 WATERGATE pulse sequence was used for solvent suppression (*80, 81*). Experiments were performed at a CRD concentration of 200 μM in 20 mM MES, 40 mM NaCl, 10 mM CaCl_2_ (pH 6.0) supplemented with 10% D_2_O and, if applicable, varying concentrations of ligands. Samples of apo DC-SIGN CRD were prepared as described for ¹H-^13^C HSQC NMR experiments. A previously published resonance assignment of DC-SIGN CRD was transferred to the nearest neighbor in a reference spectrum recorded without ligand (*62*). Unassigned, overlapping or disappearing peaks were not assigned.

CSPs induced by mutations or ligands were calculated as previously described according to **Equation 2** (*82*).

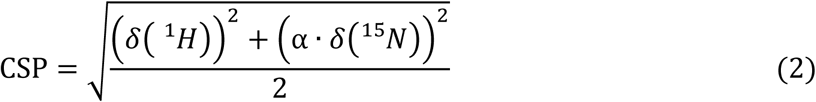

The empirical weighing factor ⍺ was set to 0.15 for all calculations.

Dissociation constants (K_D_s) were determined in titration experiments at five ligand concentrations [L]_T_ for mannose and at three for Lewis X at constant a protein concentration [P]_T_. Only residues in the fast exchange regime were used for the fitting procedure via **Equation 3** in a global two-parameter fit.

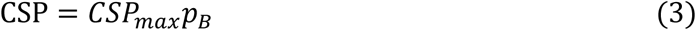

with CSP_max_ corresponding to the CSP values upon saturation and the bound protein fraction p_b_ corresponding to

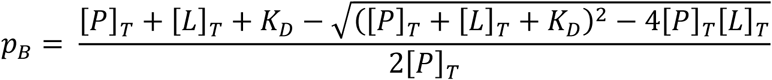

CHESPA was performed as previously described (*67*). The chemical shift vector between apo and holo wildtype DC-SIGN CRD was used as reference perturbation to calculate the vector product to determine cos(θ) of the vector between apo mutant and apo wildtype. Only residues showing CSP > 0.015 ppm were used for calculation.

#### ¹H-¹⁵N SOFAST-HMQC NMR

¹H-¹⁵N SOFAST-HMQC NMR spectra were collected with 200 increments in the nitrogen and 32 scans per increment and 1024 points in the direct dimension. The relaxation delay d1 was set to 0.3 s (*83*). Sample preparation and CSP analysis was done as described for the ^1^H-^15^N HSQC NMR experiments.

#### 19F R_2_-filtered NMR reporter displacement assay

The ^19^F R_2_-filtered NMR reporter displacement assay was adjusted for CaCl_2_ titrations with DC-SIGN ECD as previously described (*49*). Apparent transverse relaxation rates R_2,obs_ were obtained using the CPMG pulse sequence with a relaxation delay d1 of 2.0 s, acquisition time t_acq_ of 0.8 s and a CPMG frequency ν_CPMG_ of 500 Hz. DC-SIGN ECD was dialyzed four times against 25 mM HEPES, 150 mM NaCl (pH 7.4) to remove Ca²⁺ prior to measurements. Samples contained 25 μM DC-SIGN ECD in 25 mM HEPES, 150 mM NaCl (pH 7.4) supplemented with 10% D_2_O and 0.1 mM of the ManNAcF_3_ reporter molecule. TFA at a concentration of 0.1 mM served as an internal reference. 512 scans were recorded to ensure sufficient signal to noise ratios. R_2,obs_ values were obtained by fitting **Equation 4** to integrals I of the ^19^F resonance of the reporter at different relaxation times T with I_0_ as integral at T = 0 s.

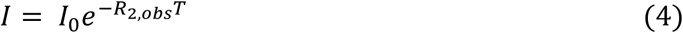

Fitted R_2,obs_ values for each CaCl_2_ concentration were used to fit EC_50_ values.

### Cryogenic electron microscopy

#### Sample preparation

The DC-SIGN ECD sample in a buffer containing 25 mM HEPES, 150 mM NaCl, 10 mM, 5mM CaCl2, pH 7.4 was thawed on ice and adjusted to a concentration of 0.02 mg mL^-1^ for vitrification. Subsequently, 3.5 µL were applied to glow-discharged holey carbon grids (Quantifoil R3/3, 300 mesh; Quantifoil Micro Tools GmbH, 1 min glow discharge), incubated for 45 s, blotted for 2 s, and plunge-frozen in liquid ethane using a Vitrobot IV device (FEI Company) operated at 4 °C and 90 % humidity.

#### Cryo-EM data acquisition and processing

Cryo-EM data were acquired at a FEI Tecnai G2 Polara microscope operating at 300 kV and equipped with a Gatan K2 Summit direct electron detector in electron-counting mode. Images were recorded at 31,000x magnification (pixel size 0.625 Å) with a total dose of 60 e⁻/Å². Automated data collection was carried out using Leginon with a defocus range of −1.5 to −2.5 μm (*84*). Gain correction, dose-weighting, motion correction, and alignment of movie frames were performed using MotionCor2 (*85*).

Subsequent data processing was carried out with cryoSPARC v4.6.0 (Fig. S6) (*86*). CTF estimation was performed with CTFFIND4 (*87*). After manual inspection of 3,574 micrographs, 3,398 micrographs were used for further analysis. Initial blob picking using a particle diameter of 350 Å and 2D classification yielded class averages that served as templates for picking particle with a diameter of 350 Å. Iterative 2D classification yielded 206,887 particles, which were extracted with a 768-pixel box size and Fourier-cropped to 192 pixels (2.5 Å/pixel). *Ab initio* reconstruction of particle images yielded two 3D reconstruction: one representing an artifactual structure (92,395 particles), and another resembling the extracellular domain of DC-SIGN (114,492 particles). The latter was subjected to a heterogeneous refinement, which removed 35,553 noisy particle images and improved the overall quality of the DC-SIGN ECD map, comprising 78,939 particles. Subsequent 2D classification and selection of high-quality 53,694 particle images, followed by *ab initio* reconstruction and non-uniform refinement, improved the definition of carbohydrate recognition domains and neck repeats in the DC-SIGN ECD structure (*88*). Finally, particle images were re-extracted using an enlarged box size of 896 pixels, Fourier-cropped to 224 pixels (2.5 Å/pixel), and further refined through *ab initio* reconstruction as well as non-uniform refinement. This resulted in a final 3D reconstruction of DC-SIGN ECD, composed of 51,929 particle images (Fig. 1J), with a low global resolution of approximately 7-8 Å, according to the Fourier shell correlation (FSC) 0.143 criterion (Fig. S7).

To evaluate the cryo-EM map of DC-SIGN ECD, a 3D structural model was generated from the human DC-SIGN ECD sequence (UniProt ID: Q9NNX6) using AlphaFold3 (*89*). The predicted atomic coordinates of DC-SIGN ECD were then rigid-body fitted into the cryo-EM map using UCSF ChimeraX (*90*). Representative views of model-to-map fit are shown in Fig. 1K, revealing differences in CRD arrangement and a reasonable fit of the neck repeats to the segmented region of the cryo-EM map.

### Mixed solvent MD simulations

#### Simulation setup

Simulations were conducted using the holo DC-SIGN in the absence of glycan ligand, retrieved from the Protein Data Bank (PDB ID: 1SL4), as the initial structural model (*52*). The system was embedded in a truncated octahedral box, ensuring a minimum solvent buffer of 10 Å between the protein surface and the box boundaries. Solvation was performed using an explicit water model supplemented with a 5% v/v concentration of phenol to create a mixed solvent environment. The AMBER ff19SB force field was employed for the parametrization of protein residues (*91*).

Initial energy minimization was carried out to relieve steric clashes and optimize the solvent configuration. The system was then gradually thermalized through a two-step protocol. The first step consisted of a 10 ps simulation under constant volume (NVT) conditions using a Berendsen thermostat. During this phase, the system was heated from 1 K to 10 K without the SHAKE algorithm, and positional restraints of 5 kcal·mol⁻¹·Å⁻² were applied to all heavy atoms of the protein to maintain structural integrity.

Subsequently, an equilibration phase was performed in two stages. The first stage involved a 5 ns simulation under isothermal-isobaric (NPT) conditions (300 K, 1 atm) using a Langevin thermostat. Hydrogen bond constraints were maintained using the SHAKE algorithm, and weak harmonic restraints of 2 kcal·mol⁻¹·Å⁻² were applied to the protein backbone atoms. The integration time step was set to 1 fs. In the second stage, a 10 ns unrestrained equilibration was conducted under the same NPT conditions, utilizing the Langevin thermostat and SHAKE constraints, with an increased integration time step of 2 fs.

Production simulations were then performed for a total duration of 1000 ns under NPT conditions at 300 K and 1 atm. Periodic boundary conditions were applied, and the SHAKE algorithm was used to constrain all bonds involving hydrogen atoms, allowing for a 2 fs integration time step. No positional restraints were imposed during the production runs.

To prevent the artificial aggregation of phenol molecules in the mixed solvent system, a dummy atom (DU) was introduced at the centroid of the aromatic ring of each phenol molecule. A repulsive Lennard-Jones potential was applied between these dummy atoms, with parameters R_min,i,j_ = 6 Å and ε_i,i_ = 0.01 kcal·mol⁻¹, as described in previous implementations of Site Identification by Ligand Competitive Saturation (SILCS) methodology.

All simulations were executed using the AMBER molecular dynamics package. Post-simulation analyses included root-mean-square deviation (RMSD) measurements, evaluation of protein-ligand interaction energies, and structural clustering to investigate conformational dynamics and binding site characteristics.

The FTMap webserver was used to compare solvent cluster of the MD-derived open state and the X-ray crystallographic structure of DC-SIGN (PDB ID: 1SL4) (*51*).

#### Determination of Water and Phenol Sites

High-occupancy regions for water and phenol were identified using Phenoltype, a VMD plugin (available upon request) based on the WatClust algorithm, with modifications to support clustering of any atom type from any cosolvent (*92*). For Water Site (WS) identification, the positions of water oxygen atoms within 5 Å of the protein surface were extracted from successive MD snapshots. Atoms within 1.4 Å of each other were clustered, and the center of mass of each cluster was defined as the WS coordinate. The occupancy of each site was calculated as the probability of finding a water molecule within a 1 Å radius of the WS coordinate, normalized to bulk solvent density. For phenol, the clustering was performed using the DU atom as the reference. Phenoltype also calculates the number of molecules per cluster and their residence times. Only sites with occupancies exceeding 10% of the total simulation time were included in subsequent analyses.

### MD simulations – Apo vs Holo DC-SIGN

#### Simulation setup

A crystal structure of the DC-SIGN CRD (PDB ID: 1K9I) was prepared for simulation of the holo state by removal of ligand, crystal waters, and all but one protein chain (*25*). To generate the apo state, the three Ca^2+^ ions were removed from the canonical binding sites. Using GROMACS 4.6, the systems were parameterized with the AMBER99SB-ILDN forcefield, solvated in TIP3P water, neutralized by addition of Cl^-^ ions, and energetically minimized using a steepest decent algorithm (emtol = 100 kJ/mol/nm, tau = 0.01) (*93–95*). Equilibrations of 100 ps in the NVT ensemble at 300 K using a Berendsen thermostat and of 150 ps in the NPT ensemble at 300 K/1 bar adding a Parrinello-Rahman barostat were carried out while putting position restraints on protein heavy atoms. Five random starting structures were selected from a 1 ns pre-run at 350 K to start one 400 ns production replica each (2000 ns total simulation time) writing solute coordinates to disk at dt = 1 ps. A leap-frog integrator was used with a 2 fs time step. Periodic boundary conditions were applied in x-, y-, and z-direction.

#### Mutual information analysis

Backbone and side chain dihedral angles were extracted from MD trajectories using MDTraj (*96*). Normalized mutual information (NMI) values were calculated between all pairs of φ-ψ vs. φ-ψ (2D2D), φ-ψ vs. χ (2D1D), and χ vs. χ (1D1D) distributions using a custom Cython implementation (*97*). The input distributions were discretized using 90 bins per dimension (Fig. S11). The obtained angle-wise NMI contributions were projected onto residue-wise contributions and not normalized but the resulting matrix was centered via 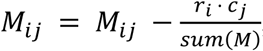, with row sums r and column sums c.

### Alignment and sequence analysis

Unique human structures of C-type lectins deposited in the PDB (38 total) were chosen for structural and sequence alignment using the PyMol and the MAFFT software (*98*). Sequence logos were created using the WebLogo 3 software (*99*).

## Acknowledgments

We would like to thank the NMR facility at the Department of Pharmaceutical Sciences, Faculty of Life Sciences, University of Vienna, for providing critical support and access to instrumentation for this work. CPM thanks the German Academic Exchange Service (DAAD) for financial support.

## Author contributions

Conceptualization: CR, BK, CPM, JL

Methodology: JL, CPM, BK, JOKJ, PS

Investigation: JL and MB conducted protein expression, NMR spectroscopy and biochemical assays with contributions from HF and GS, MDG and CPM conducted MD simulations, JOKJ analyzed MD simulations with support from EG, AB conducted cryo-EM experiments

Formal analysis: JL, MDG, CPM, JOKJ, AB

Writing – original draft: JL and CR

Writing – review and editing: JL, CR, MB, CPM, JOKJ, BK, AB, PS

Resources: CR, BK, PS, CPM

Supervision: CR, BK, PS, CPM

Funding acquisition: CR, BK, PS, MB

