## Supplementary Information for "A cryptic pocket allosterically modulates oligosaccharide binding to DC-SIGN"

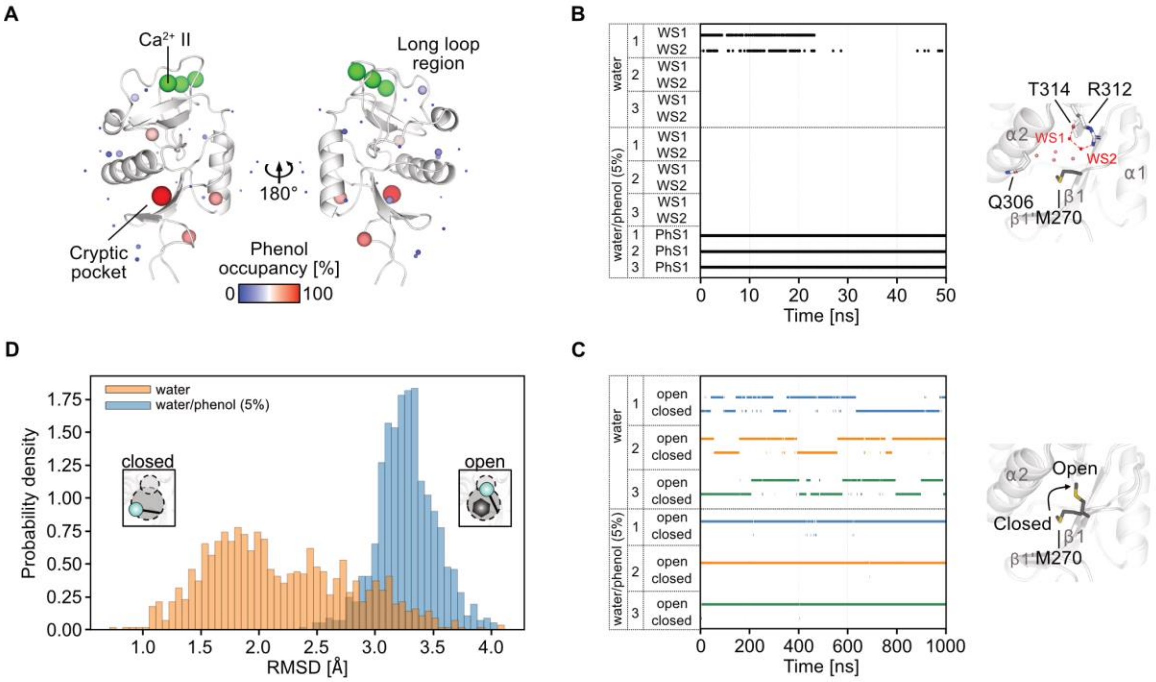


**Fig. S1. Mixed solvent MD simulations with phenol uncover a cryptic pocket. (A)** Phenol occupancy clusters on the DC-SIGN CRD surface over the course of the simulations. Twenty-seven clusters appearing in ≥10% of the total simulation time are shown as spheres, scaled and colored by relative occupancy. The cryptic pocket is identified as the most frequently occupied site. **(B)** Solvent structure analysis of the cryptic pocket during the final 50 ns of each 1 µs simulation replicate (1–3). Hotspots for water molecules WS1 and WS2, located in the upper cavity near T314 and R312, can be displaced by M270 side-chain rotation in both water (intermittently) and 5% phenol simulations (persistently). When phenol is present at the phenol hotspot representing the cryptic pocket (PhS1), it occupies the cryptic pocket and stabilizes the open conformation, preventing reformation of WS1 and WS2. **(C)** Cluster-based time-series analysis of the conformation of residue M270 relative to its conformation in PDB ID: 1SL4. All frames across the full simulation time with a side chain RMSD ≤ 2 Å were labeled as "closed," while those with RMSD ≥ 2.5 Å were tagged as "open." Intermediate values were considered negligible for this specific analysis. In water, M270 alternates between open and closed conformations, while in phenol-containing simulations, phenol binding stabilizes the open state for the majority of the trajectory. **(D)** Distribution of M270 side chain RMSD values relative to the closed conformation across all frames and replicates. The probability density plot summarizes the same data used in **(C)**, showing a bimodal distribution in water (orange) and a shift toward the open state in 5% phenol (blue). RMSD values were binned and normalized to produce probability densities, highlighting the phenol-induced stabilization of the open conformation.

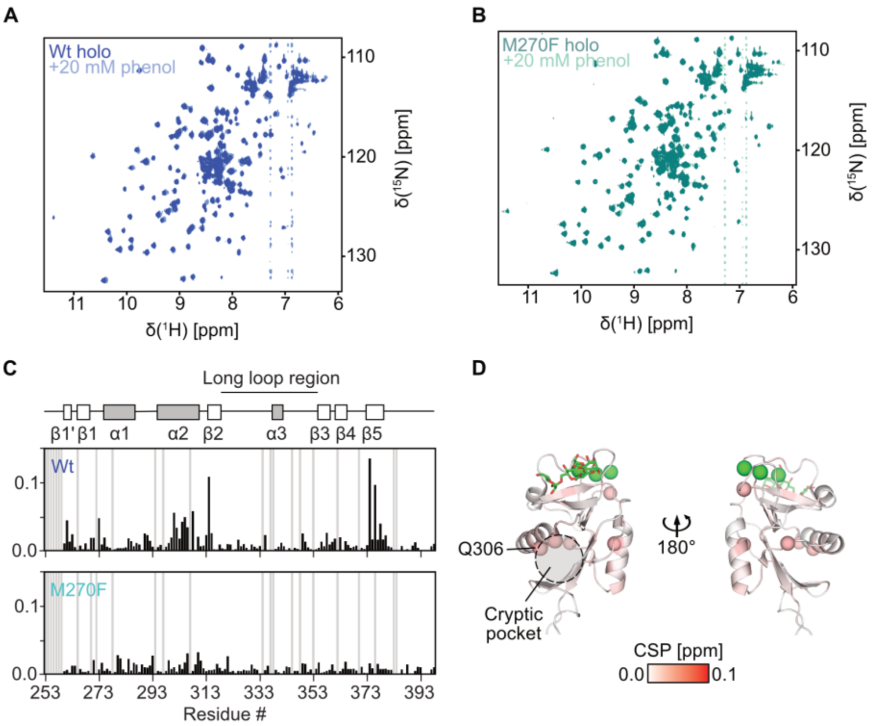


**Fig. S2. The M270F mutation blocks DC-SIGN from interacting with phenol. (A)** and **(B)** $^{1}$H-$^{15}$N HSQC NMR spectra of holo DC-SIGN CRD wildtype and M270F in the absence or presence of phenol. **(C)** Comparison of CSPs maps of the wildtype and M270F interacting with 20 mM phenol, show drastically reduced CSPs for the mutant. **(D)** Mapping of CSPs observed addition of 20 mM phenol to holo DC-SIGN CRD M270F confirm inhibition of phenol binding by the mutant. Cα atoms of residues experiencing a CSP > 0.025 ppm are shown as spheres.

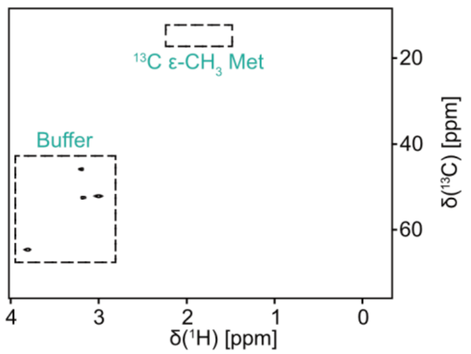


**Fig. S3. $^{1}$H-$^{13}$C HSQC NMR spectrum without $^{13}$C ε-methyl methionine labeled DC-SIGN CRD wildtype.** Spectra of samples containing only buffer (20 mM MES, 40 mM NaCl, 10 mM CaCl2, pH 6.0) without labeled protein served to locate buffer peaks and exclude scrambling of the $^{13}$C ε-methyl methionine during protein expression.

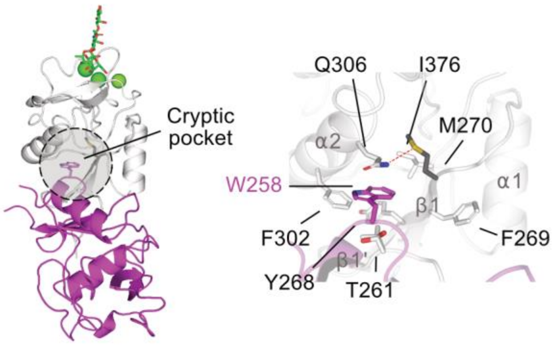


**Fig. S4. The open cryptic site in the X-ray crystallographic structure of DC-SIGN CRD wildtype.** Residue W258 from a symmetry mate (pink cartoon) in the X-ray crystallographic structure of DC-SIGN CRD in complex with lacto-N-fucopentaose III (white cartoon, PDB: 1SL5) opens the cryptic pocket. M270 rotates into the upper cavity and is additionally stabilized by a predicted hydrogen bond (red dotted line) to residue Q306 in α-helix 2.

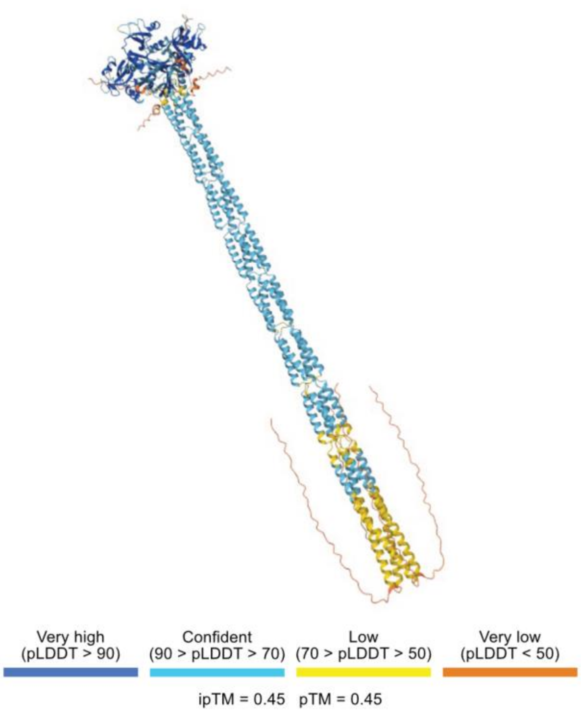


**Fig. S5. AlphaFold3-predicted structure of the human DC-SIGN ECD.** The model was generated using the protein sequence of human DC-SIGN ECD (Uniprot ID: Q9NNX6). Confidence levels of the predicted structure are color-coded as indicated by the scale bar.

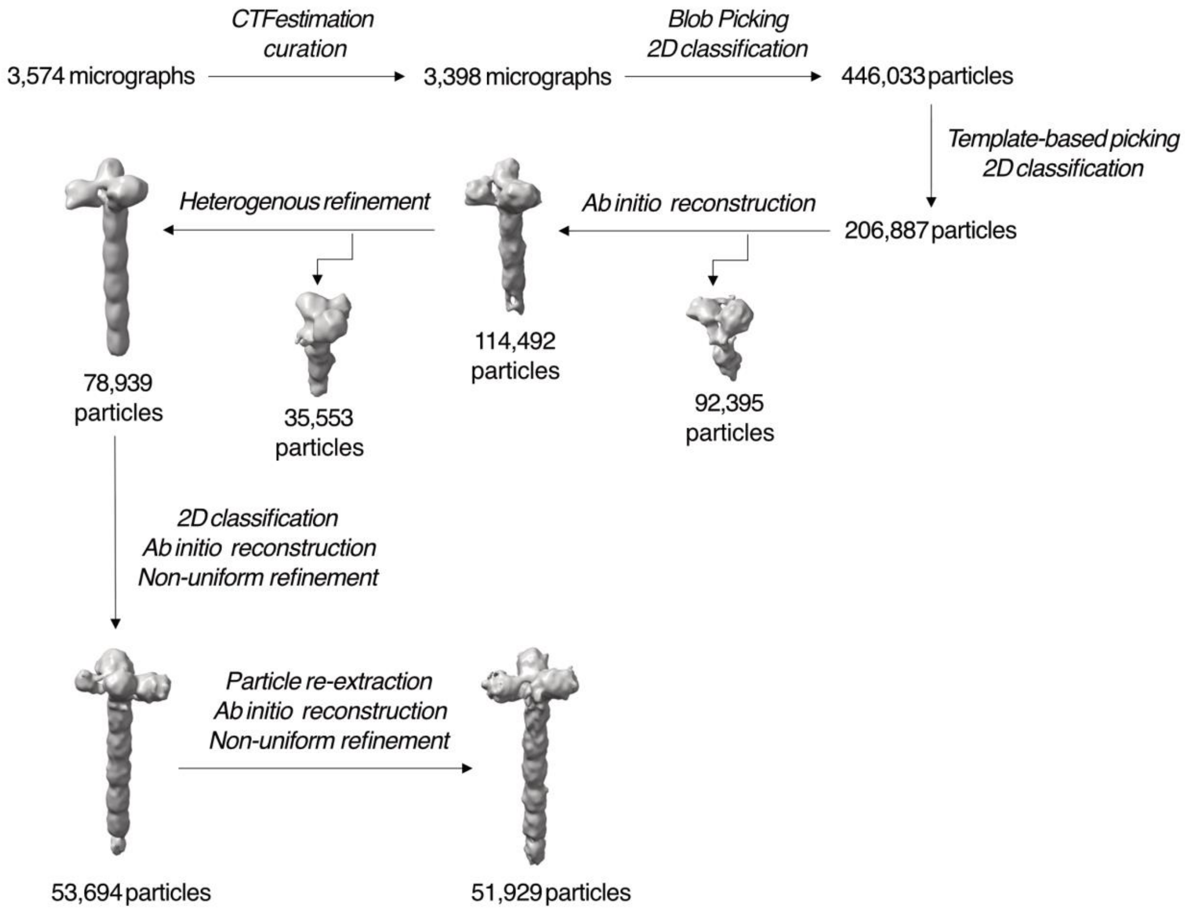


**Fig. S6. Cryo-EM refinement sorting scheme of the DC-SIGN ECD dataset.** After CTF estimation and manual micrograph inspection, initial blob picking and 2D classification were performed. The resulting 2D class averages served as templates for particle picking. Iterative 2D classification yielded 206,887 particle images, which were extracted with a box size of 768 pixels and subsequently Fourier-cropped to 192 pixels (2.5 Å/pixel). Ab initio reconstruction of these particle images revealed a 3D reconstruction resembling the expected structure of DC-SIGN ECD. This 3D reconstruction was subjected to heterogeneous refinement, which removed noisy particles and improved the overall DC-SIGN ECD structure, consisting of 78,939 particle images. Subsequent 2D classification and selection of high-quality particle images, followed by ab initio reconstruction and non-uniform refinement, improved the definition of the CRDs and neck repeats in the DC-SIGN ECD structure. Finally, these particle images were then re-extracted with an increased box size of 896 pixels, Fourier-cropped to a box size of 224 pixels (2.5 Å/pixel) and subjected once more to ab initio reconstruction and non-uniform refinement. This resulted in a final 3D reconstruction of DC-SIGN ECD at a global resolution of 7-8 Å (FSC 0.143 criterion).

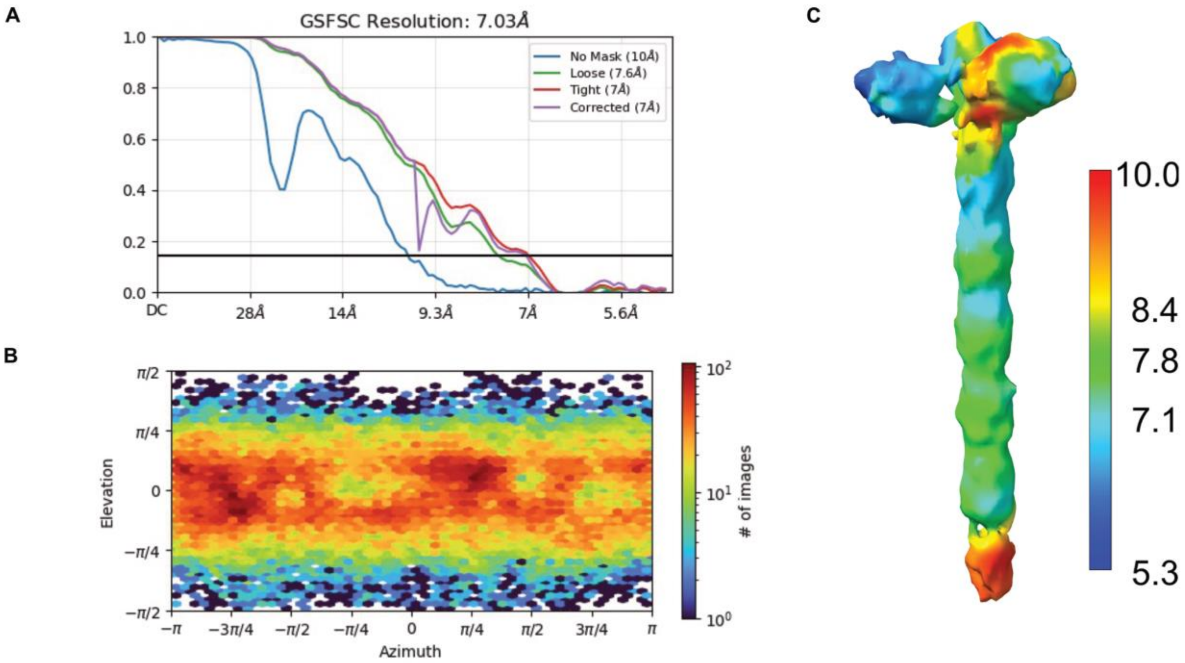


**Fig. S7. Global resolution estimation of cryo EM maps of DC-SIGN ECD. (A)** The global resolution estimation, according to the FSC 0.143 criterion, was performed in cryoSPARC. **(B)** Projection direction distribution from non-uniform refinement in cryoSPARC, showing a preference for side views and underrepresentation of top views. **(C)** Final cryo-EM map colored by its local resolution estimation ranging from 5.3 to 10.0 Å as indicated by the colored scale bar.

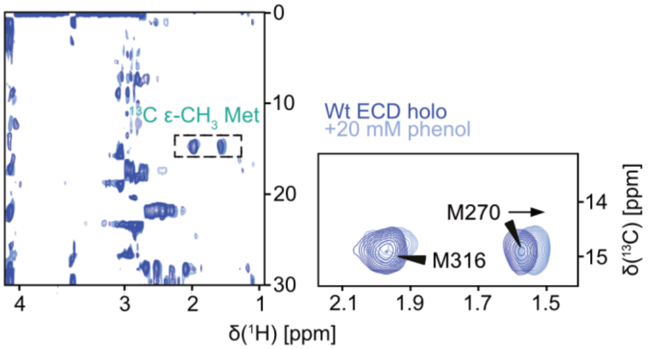


**Fig. S8. The cryptic pocket is accessible in DC-SIGN ECD in solution.** $^{1}$H-13C TROSY NMR spectra of $^{13}$C ε-methyl methionine labeled DC-SIGN ECD in the presence or absence of 20 mM phenol. The M270 resonance shows a clear shift similar to what is observed in the CRD (Fig. 1G), indicating that the cryptic pocket is accessible in the tetrameric ECD.

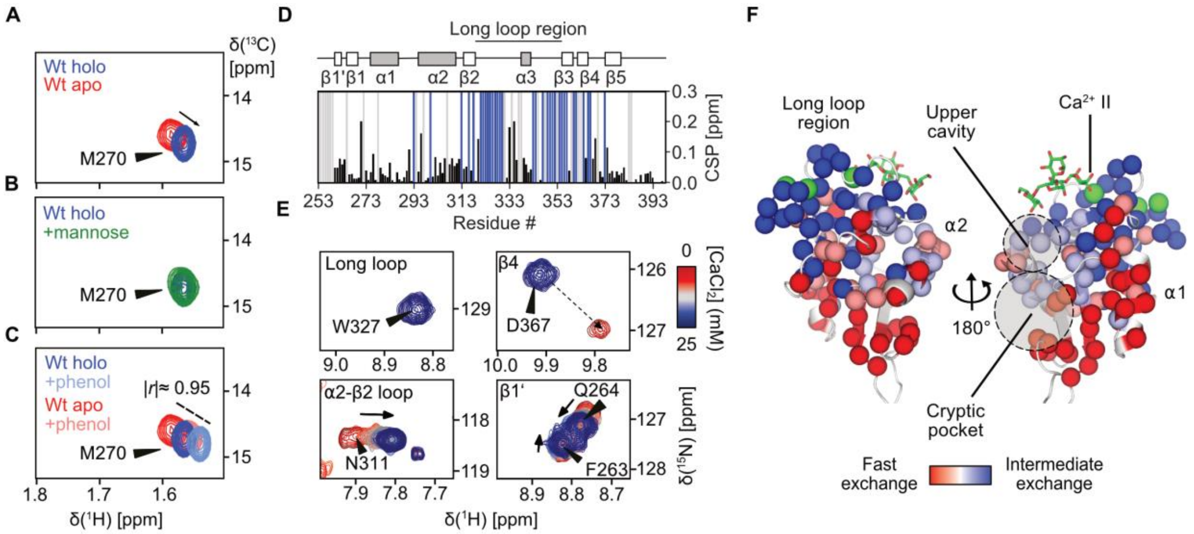


**Fig. S9. $Ca^{2+}$ binding induces conformational change throughout the CRD. (A)** Removal of $Ca^{2+}$ from $^{13}C$ ε-methyl methionine-labeled DC-SIGN CRD shifts the resonance of M270 indicating conformational change at the cryptic site. **(B)** Addition of 20 mM mannose to holo DC-SIGN CRD does not induce shift in M270. **(C)** Addition of 20 mM phenol to apo DC-SIGN shifts M270 along the same trajectory towards the holo state. Pearson correlation of the chemical shift position indicates linear correlation of the holo, and the phenol bound state. **(D)** CSPs induced by addition of 25 mM $CaCl_2$ to $^{15}N$-labeled apo DC-SIGN CRD wildtype in $^1H$-$^{15}N$ HSQC NMR experiments reveals global conformational change throughout the CRD. **(E)** Residues in different regions of the CRD show different exchange regimes upon titration of $Ca^{2+}$. **(F)** Qualitative mapping of exchange regimes as observed in $Ca^{2+}$ titrations reveal α-helix 2 to be in fast-to-intermediate exchange, while the long loop region is in intermediate exchange and more distal regions, including the cryptic site, are in fast exchange.

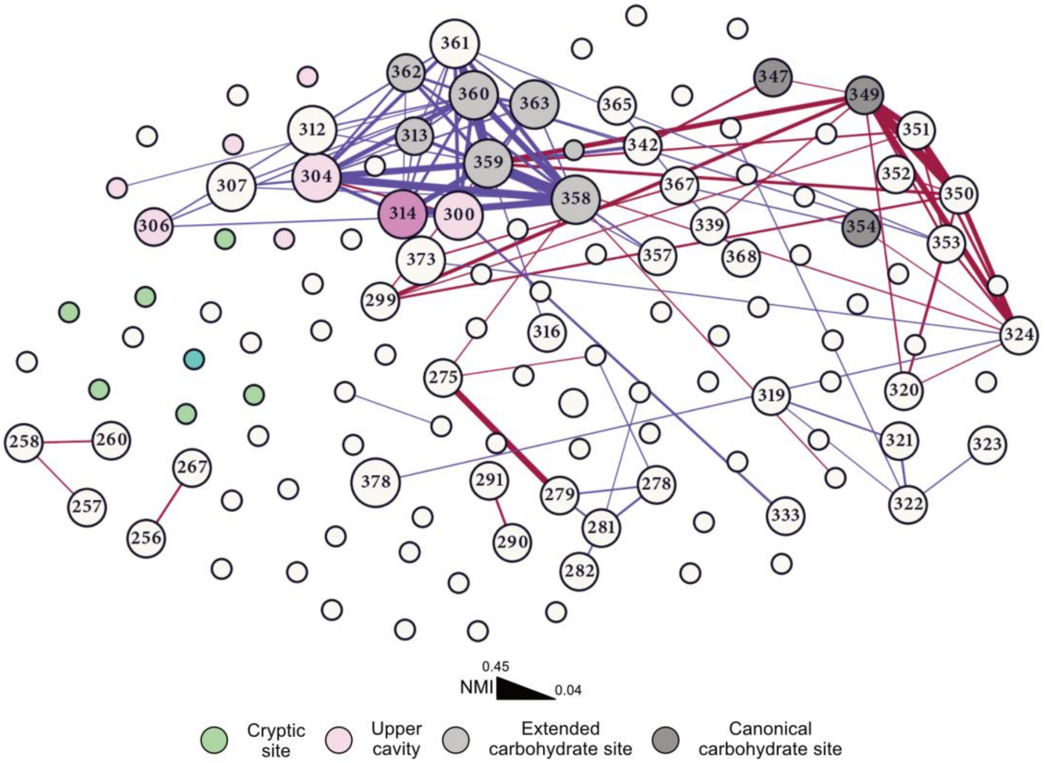


**Fig. S10. $Ca^{2+}$ binding shifts the connectivity of the DC-SIGN CRD.** Network representation of the difference in NMI values computed for the holo and apo state (holo - apo) of DC-SIGN CRD. Blue and red edges illustrate higher NMI values in the holo and apo state, respectively. Scaling and coloring of nodes and edges as described for Fig. 2A.

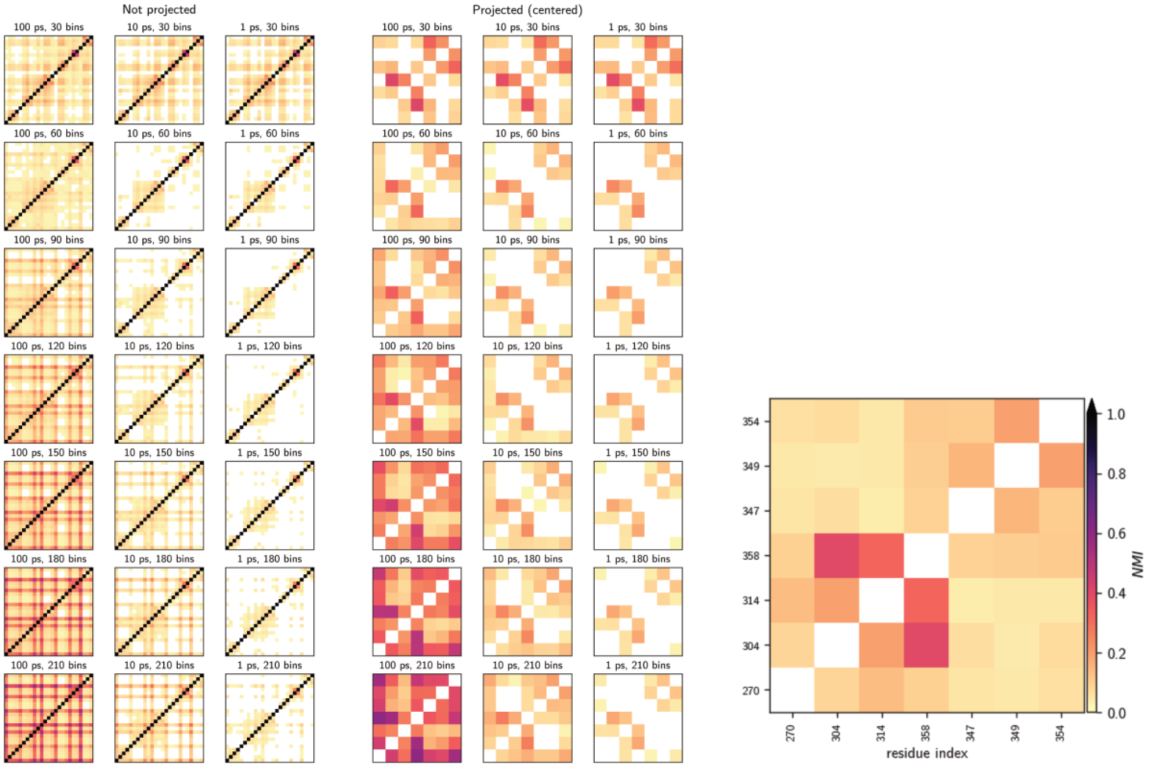


**Fig. S 11. Effect of striding and binning on computed NMI values.** For all pairs of dihedral angle distributions in residues 270, 304, 314, 347,349, 354, and 358 we screened trajectory strides of 1, 10, and 100 ps in combination with 30 to 210 histogram bins when computing NMI values and plotted the original (left) and residue –wise projected (right) NMI matrices. In order to avoid poor resolution as well as noisy estimates, we settled on using 1 ps/90 bins for the full analysis.

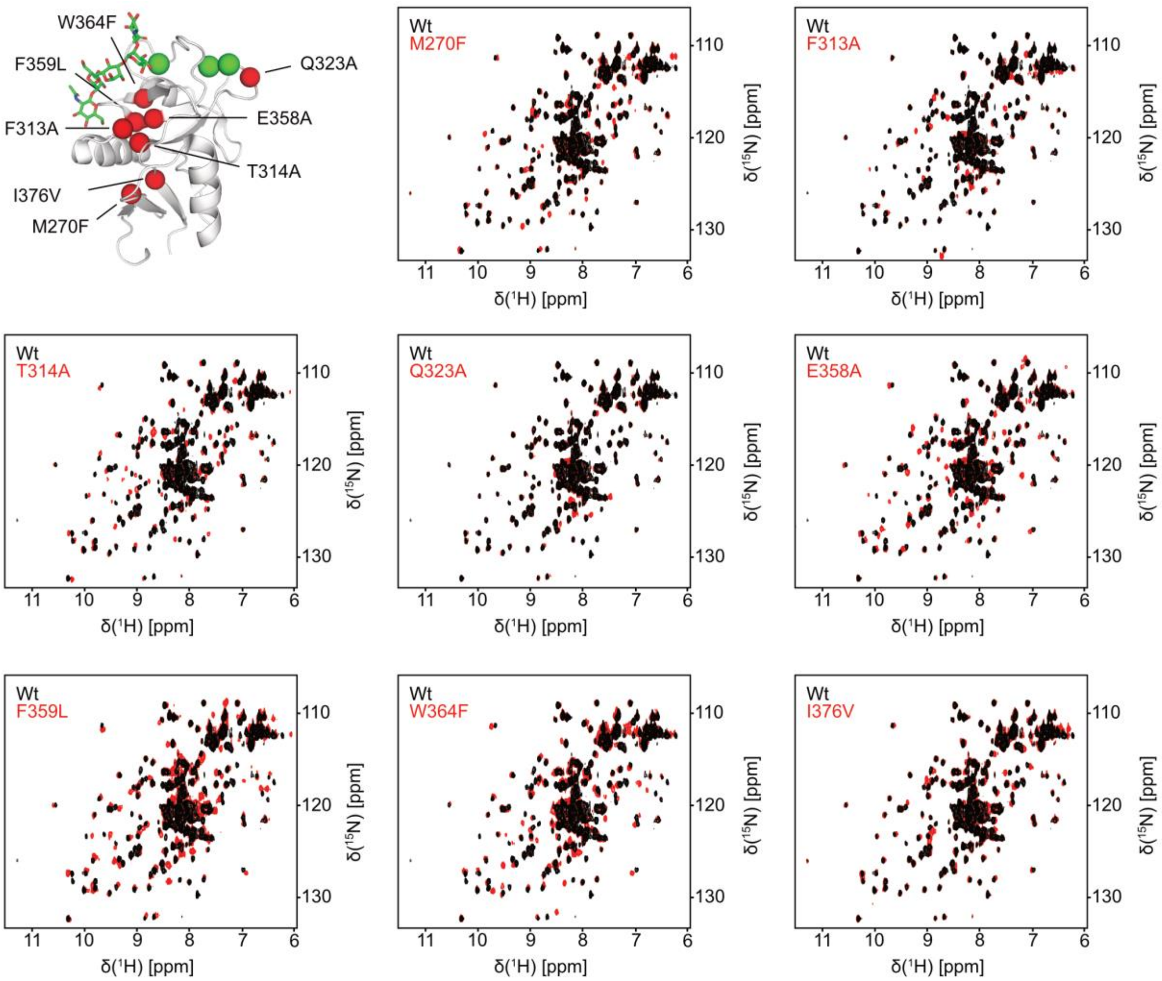


**Fig. S12. Point mutations in DC-SIGN yield folded CRDs.** Locations of point mutation mapped to the X-ray crystallographic structure of DC-SIGN CRD (PDB ID: 1K9I). Red spheres indicate Cα atoms of positions selected for mutation in the DC-SIGN CRD. $^{1}H$-$^{15}N$ HSQC NMR spectra of holo DC-SIGN CRD wildtype and mutants show dispersed spectra with significant chemical shift differences to the wildtype protein.

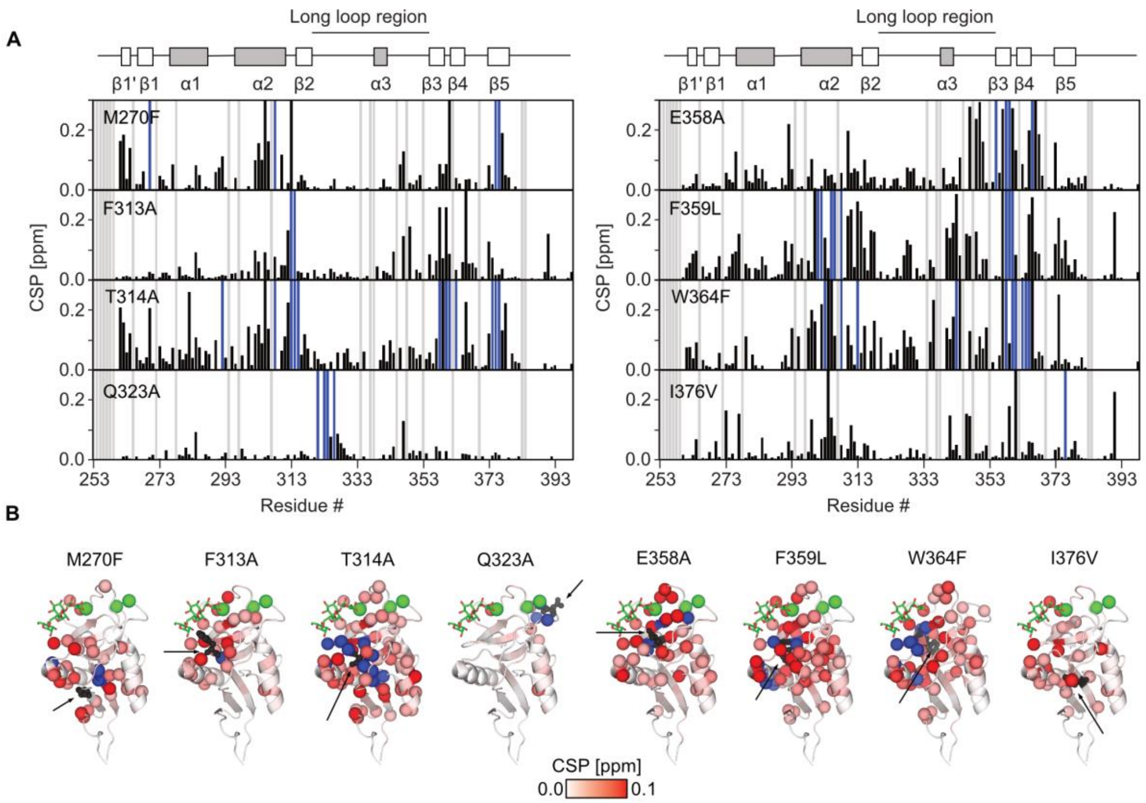


**Fig. S13. Mutation of hub residues induce chemical shift perturbations distal to the mutation site. (A)** CSP maps illustrating chemical shift changes induced by point mutations relative to the wildtype in $^{1}H$-$^{15}N$ HSQC NMR spectra. Grey bars indicate unassigned residues. Blue bars indicate resonances of residues that experience severe line broadening or where assignment could not be unambiguously transferred due to large CSPs. **(B)** CSPs induced by point mutations mapped to the structure of DC-SIGN CRD. Cα atoms of residues experiencing a CSP > 0.025 ppm are shown as spheres. Blue spheres indicate resonances of residues that experience severe line broadening or where assignment could not be unambiguously transferred due to large CSPs. Mutated residues are shown as black spheres.

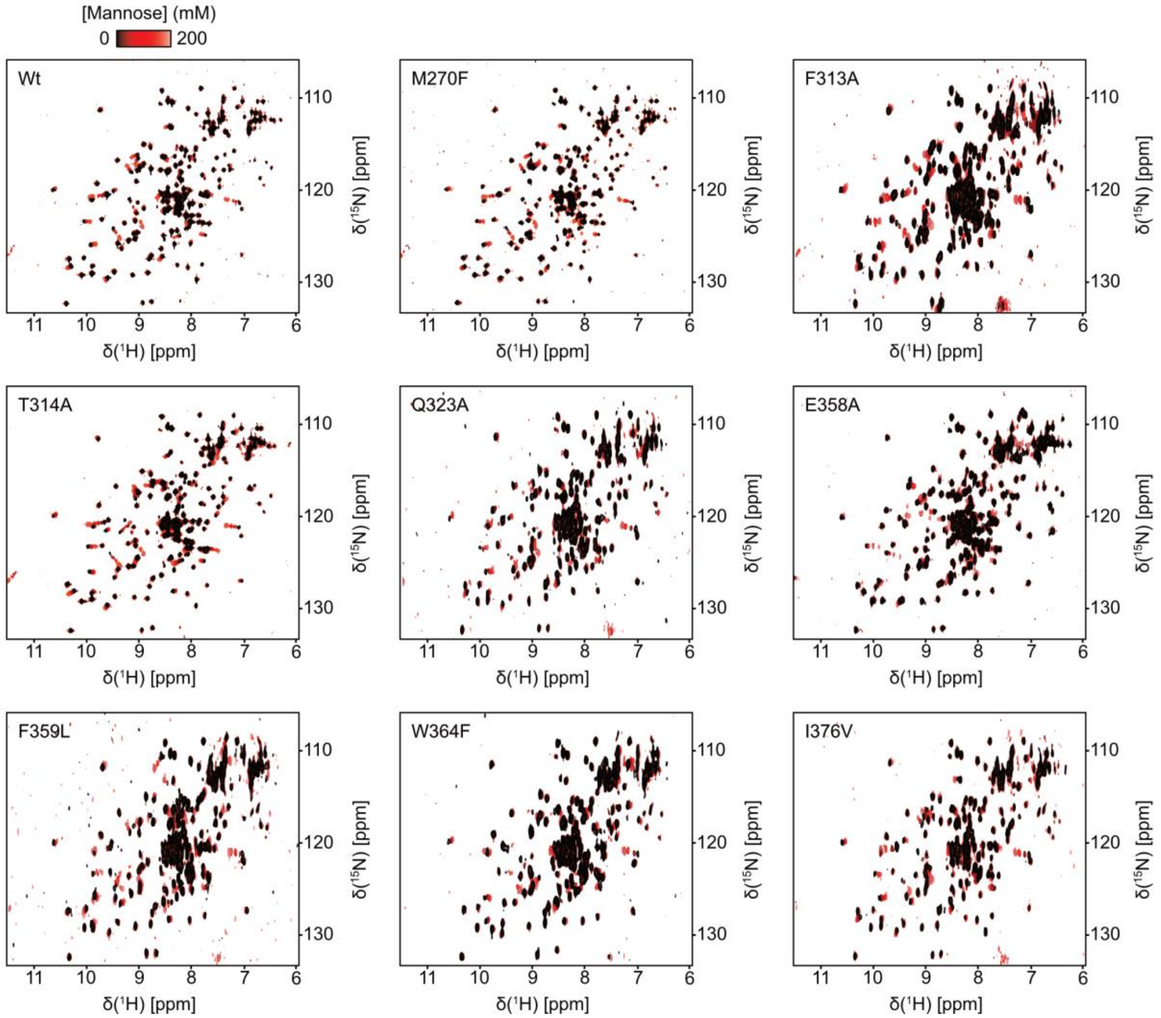


**Fig. S14. The mutant CRDs bind mannose.** $^{1}$H-$^{15}$N HSQC (Wt, M270F, T314A, E358A) and SOFAST-HMQC (F313A, Q323A, F359L, W364F, I376V) NMR spectra of holo DC-SIGN CRD wildtype and mutants at varying concentrations of mannose. All proteins show characteristic CSPs upon addition of mannose, indicating correct folding and functionality of the CRDs.

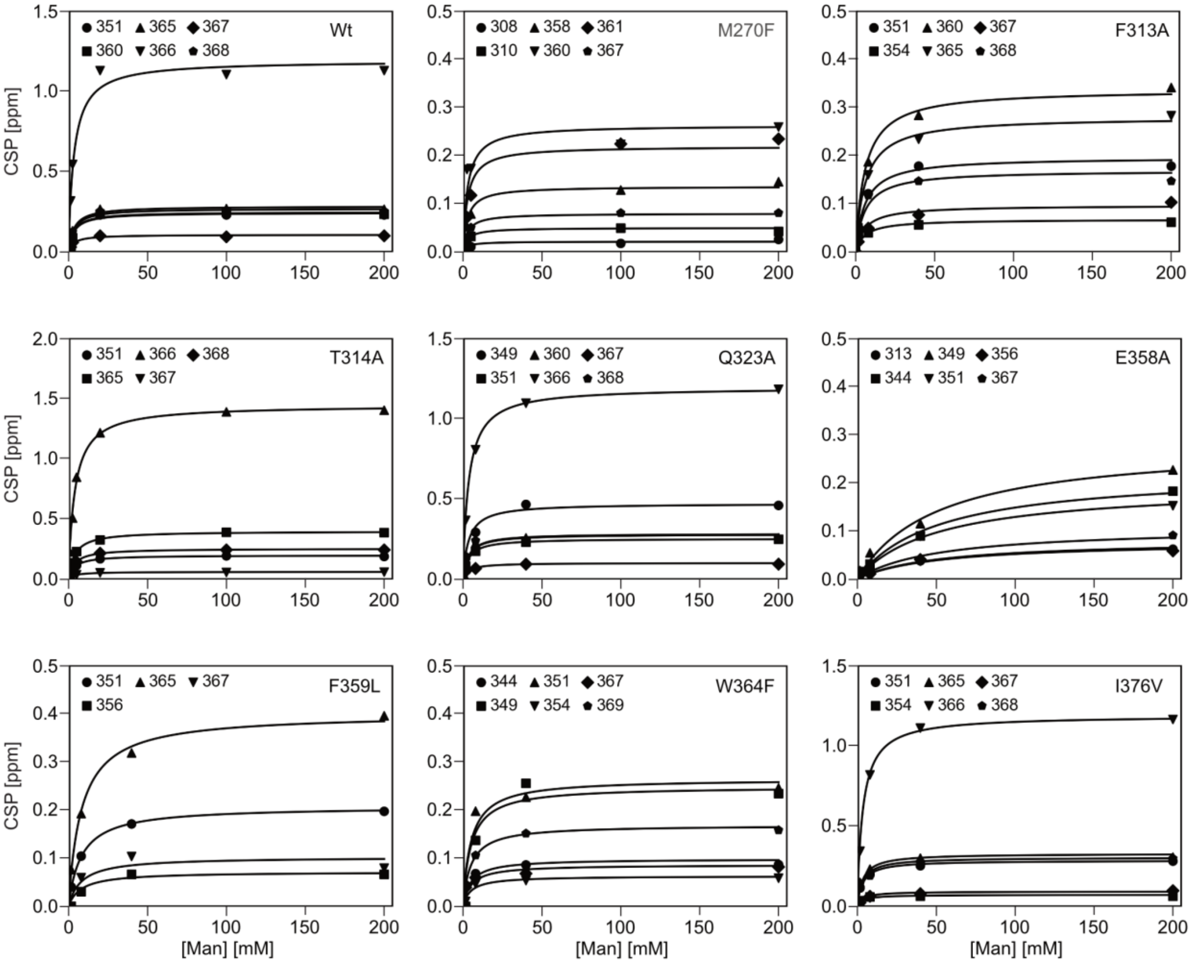


**Fig. S15. Affinities of wildtype and mutant CRDs interacting with mannose.** Global fitting of CSP trajectories reveals similar mannose affinities for the wildtype ($K_D$ = 3.3 ± 0.3 mM) and for the M270F ($K_D$ = 2.9 ± 0.4 mM), T314A ($K_D$ = 3.9 ± 0.1 mM), Q323A ($K_D$ = 3.5 ± 0.2 mM), and I376V ($K_D$ = 3.4 ± 0.2 mM) mutants, while mutations of hub residues F313A ($K_D$ = 6.0 ± 0.5 mM), E358A ($K_D$ = 51.1 ± 4.9 mM), F359L ($K_D$ = 8.2 ± 0.8 mM) and W364F ($K_D$ = 5.0 ± 0.9 mM) show reduced affinity.

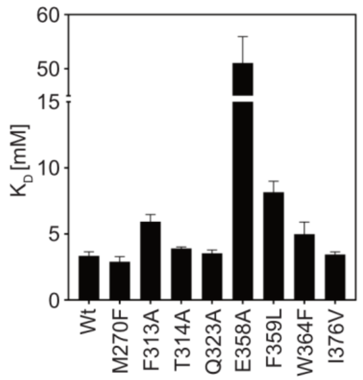


**Fig. S16. Mutation of hub residues differentially modulate mannose affinity.** Comparison of fitted mannose affinities from $^1H$-$^{15}N$ HSQC NMR titrations (see also Fig. S15).

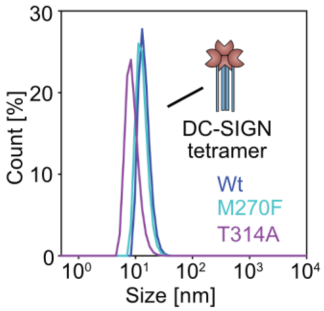


**Fig. S17. The mutant proteins form oligomers in solution.** DLS measurements of the ECD proteins in 25 mM HEPES, 150 mM NaCl, 10 mM CaCl2, pH 7.4. Inflection points reveal no significant changes in tetramerization properties.

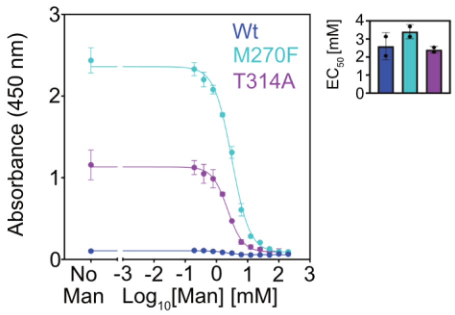


**Fig. S18. Inhibition of HRP binding by mannose.** Binding of HRP at constant concentration to the ECD proteins at varying mannose concentrations, reveal marginal changes in the mannose $IC_{50}$ for the M270F ($IC_{50}$ = 3.1 mM, hill slope = -1.5) and T314A ($IC_{50}$ = 2.1 mM, hill slope = -2.1) mutations compared to the wildtype ($IC_{50}$ = 2.3 mM, hill slope = -1.8).

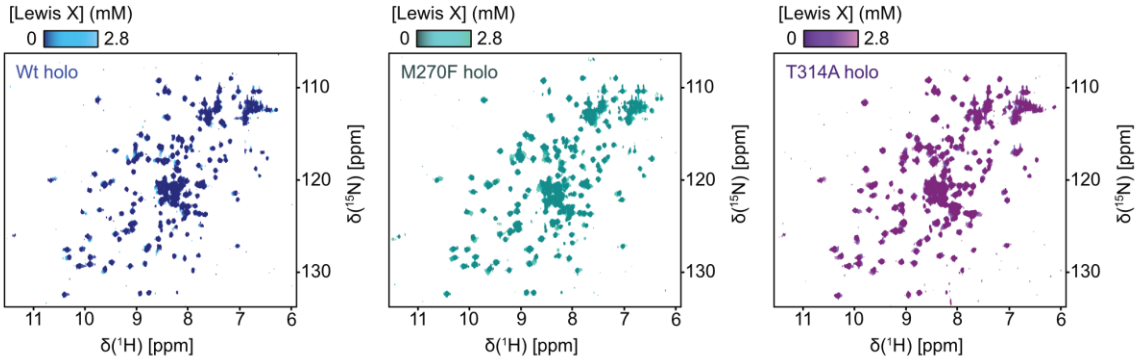


**Fig. S19. The mutant CRDs bind Lewis X.** $^{1}H$-$^{15}N$ HSQC NMR spectra of holo DC-SIGN CRD wildtype and mutants at varying concentrations of the Lewis X trisaccharide. All proteins show characteristic CSPs upon addition of Lewis X.

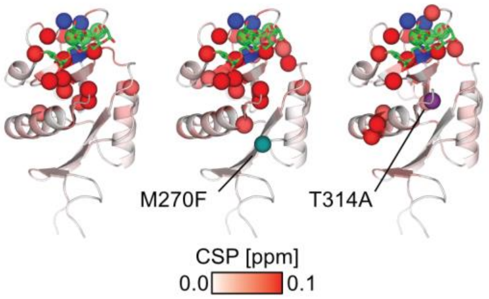


**Fig. S20. Lewis X induces CSPs at the canonical and extended carbohydrate binding site.** Mapping of CSPs induced by the addition of 2.8 mM Lewis X to the structure of DC-SIGN CRD (PDB ID: 1SL4), suggest the ligand to interact with the canonical and the extended carbohydrate binding site. Cα atoms of residues experiencing a CSP > 0.025 ppm are shown as spheres. Blue spheres indicate resonances of residues that experience severe line broadening or where assignment could not be unambiguously transferred due to large CSPs. M270F and T314A mutation sites are shown as dark cyan and purple spheres, respectively.

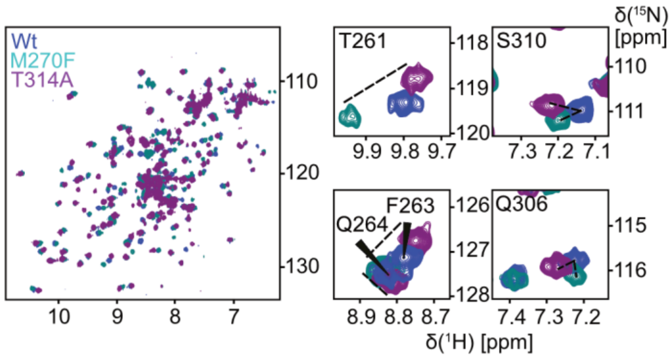


**Fig. S21. Mutations M270F and T314A induce linear and non-linear chemical shift changes.** Comparison of $^{1}$H-$^{15}$N HSQC NMR spectra of holo DC-SIGN CRD wildtype, M270F and T314A. Resonances of residues T261, F263 and Q264 are shown as example for linear displacement of chemical shifts upon mutation. Resonances of residues S310 and Q306 are shown as examples for non-linear behavior.

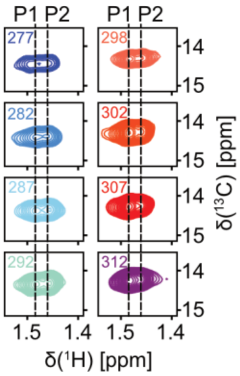


**Fig. S22. Temperature-dependent chemical shift changes of the M270 resonance in apo DC-SIGN T314A.** $^{1}H$-$^{13}C$ HSQC NMR spectra of $^{13}C$ ε-methyl methionine-labeled apo DC-SIGN CRD T314A at different temperatures. Only the resonance of M270 is shown. With increasing temperature, state P1 becomes more populated, indicating an entropically driven transition from the holo-like state P2.

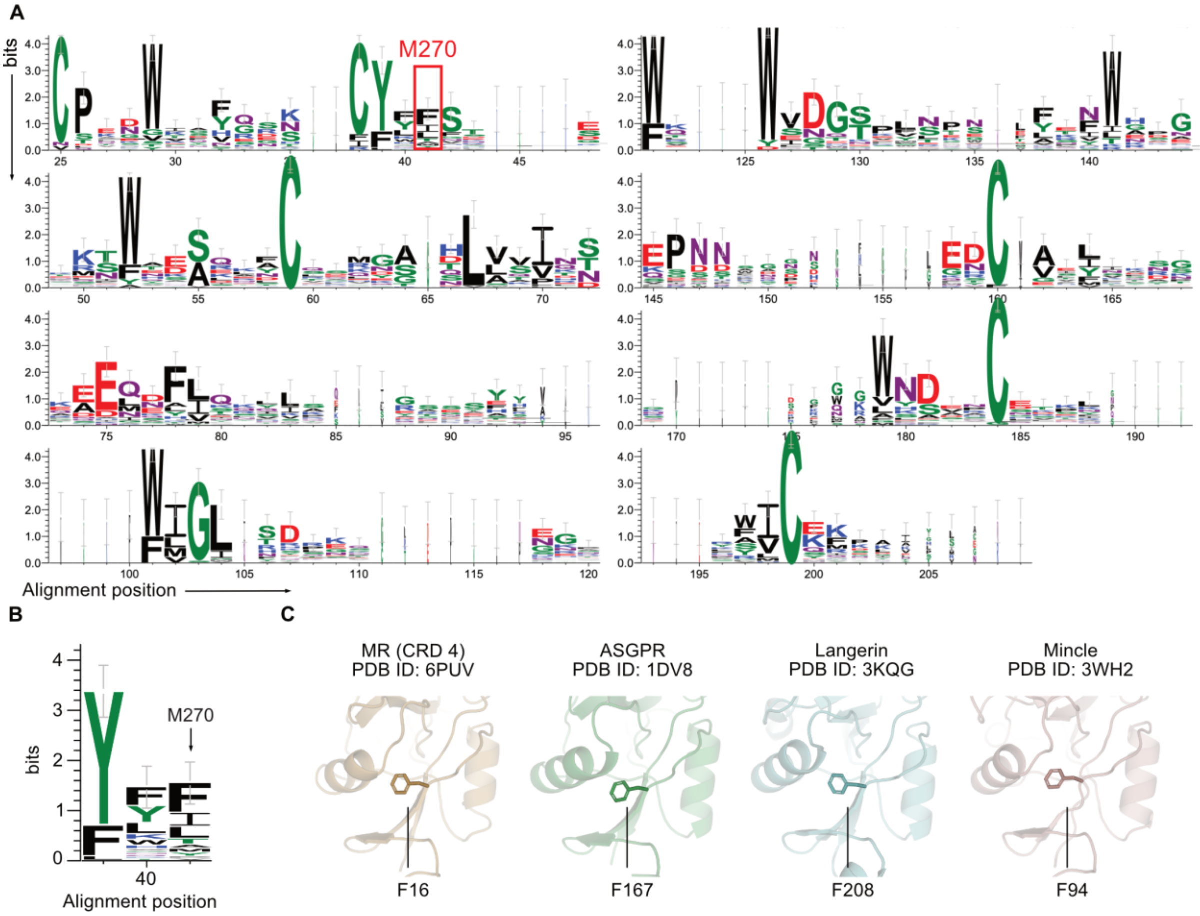


**Fig. S23. Phenylalanine is evolutionary preferred in position of M270. (A)** Amino acid sequence logo generated from an alignment of 38 human C-type lectin domain structures. The alignment position 41 corresponding M270 in DC-SIGN is highlighted in red. Letter height denotes information content. Alignment positions before position 25 showed low conservation and were left out for clarity. **(B)** Inset of sequence logo, showing phenylalanine as the preferred amino acid in position of M270. **(C)** Example C-type lectins expressing phenylalanine in position of M270.

**Tabel S1. Selected hydrogen bond populations of apo and holo DC-SIGN.**

| | | holo | apo | |
|---|---|---|---|---|
| **Donor[a]** | **Acceptor[a]** | **Occupancy** | **Occupancy** | **ΔHB[b]** |
| N322s | T326m | 18.57% | 10.00% | 8.57% |
| S360m | G363m | 15.85% | 9.80% | 6.05% |
| N344s | S360m | 5.84% | | 5.84% |
| S360s | E358s | 4.79% | | 4.79% |
| V292m | I376m | 37.73% | 34.19% | 3.54% |
| K379s | V287m | 46.35% | 43.10% | 3.25% |
| T314s | S307s | 3.25% | | 3.25% |
| F374m | F313m | 2.95% | | 2.95% |
| W315m | F374m | 75.49% | 72.65% | 2.84% |
| A382m | P255m | 21.82% | 19.04% | 2.78% |
| D320m | D355m | 25.92% | 23.41% | 2.51% |
| T261m | Y268m | 45.24% | 42.85% | 2.39% |
| G363m | S360m | 2.39% | | 2.39% |
| R345m | N344s | 2.29% | | 2.29% |
| N266m | F263m | 21.73% | 19.52% | 2.21% |
| Y342m | F339m | 5.96% | 3.85% | 2.11% |
| R275s | D279m | 40.64% | 38.55% | 2.09% |
| R275s | D279s | 30.87% | 28.79% | 2.08% |
| R345s | N344s | 1.09% | 3.10% | -2.01% |
| K379m | C267m | 36.90% | 39.14% | -2.24% |
| N362m | S360s | 9.82% | 12.28% | -2.46% |
| F302m | E298m | 34.03% | 36.49% | -2.46% |
| N350m | D366s | 67.66% | 70.30% | -2.64% |
| F359m | T314m | 80.85% | 83.53% | -2.68% |
| S307s | L303m | 61.38% | 64.08% | -2.70% |
| Q341m | Q341s | 12.04% | 14.90% | -2.86% |
| S308s | Q304m | 36.55% | 39.59% | -3.04% |
| Q304s | G361m | 32.29% | 35.72% | -3.43% |
| T314m | F359m | 26.58% | 30.05% | -3.47% |
| Q300s | Q304s | 22.93% | 27.14% | -4.21% |
| S319s | D355m | 15.67% | 20.32% | -4.65% |
| W364s | Q300s | 40.17% | 44.98% | -4.81% |

[a] s = sidechain, m = main chain

[b] Difference between populations of hydrogen bonds observed in simulations of holo - apo state of DC-SIGN CRD.
